# Suspended Tissue Engineering with Assemblable Microfluidics (STEAM)

**DOI:** 10.1101/2025.10.28.684690

**Authors:** Jamison M. Whitten, Amanda J. Haack, Liam A. Knudsen, Ella E. Bouker, Asha R. Viswanathan, Lauren G. Brown, Dahna A. Kim, Laura A. Milton, Ariel Lin, Alex Georgiou, Emme A. Schumacher, M. Yunos Alizai, Yi-Chin Toh, Jean Berthier, Cole A. DeForest, Nathan J. Sniadecki, Ashleigh B. Theberge, Erwin Berthier

## Abstract

Suspended tissue culture systems enable cellular responses to mechanical forces critical for tissue development and function. Tissues develop in complex environments containing both mechanical and chemical cues that vary spatiotemporally; thus modeling both of these physiochemistries *in vitro* through integration of spatial patterning with mechanical manipulation simultaneously is an important aspect in microphysiological tissue modeling which has yet to be achieved. Here, we introduce Suspended Tissue Engineering with Assemblable Microfluidics (STEAM), a modular tissue fabrication platform that allows for spatially heterogeneous suspended tissue architectures. With STEAM, we achieve tissue constructs with multiple regions through the addition of capillary pinning features to control hydrogel precursor flow. STEAM tissues can easily be moved from patterning setup to well-plate to microscope slide, which also enables stacking of separately generated layers. Mechanical manipulation post-fabrication is also possible via static stretching, where cell-embedded 3D tissues can be stretched farther apart to induce strain along an axis. To demonstrate the utility of post fabrication strain ability, we showed that myotube alignment increases when strain is applied to STEAM generated engineered muscle tissue containing mouse myoblasts. Finally, by modifying the channel geometry of the fluidic-based patterning rails, we generate complex nonplanar suspended tissues. STEAM leverages microfluidic principles to generate suspended tissues that integrate patterning precision, mechanical functionality, and experimental versatility, providing a suite of construct combinations for modeling tissue behaviors from the interplay of spatial organization and mechanical forces.

## INTRODUCTION

Mechanical forces govern tissue development, homeostasis, and disease progression, with mechanical dysfunction underlying cardiovascular disease, muscle injury, and fibrotic disorders.^1^ Tissue function emerges at the intersection of mechanical forces and spatial interfaces between heterogeneous tissue regions, requiring *in vitro* platforms that integrate both spatial control and mechanical manipulation.^2,3^ Suspended tissue culture systems, in which constructs are anchored between flexible or rigid posts or pillars, have proven effective for generating mechanically functional tissues by enabling cells embedded in three-dimensional (3D) extracellular matrix (ECM) to generate physiological tension and respond to applied forces.^4,5^ These platforms have proven invaluable for cardiac tissue maturation,^6–8^ skeletal muscle engineering,^9,10^ and mechanobiology studies,^11–13^ establishing that tissues cultured under tension develop superior contractile function, alignment, and phenotypic maturity compared to unstrained constructs.

While suspended tissue platforms have integrated multiple functionalities (e.g., combining perfusion with mechanical strain in organ-chips^14^ or force measurement with electrical stimulation^9,15–17^), the combination of spatial patterning (e.g. controlling the arrangement of materials), post-fabrication mechanical manipulation, and tissue portability remains challenging. Spatial heterogeneity is critical for modeling disease-healthy border zones (e.g., myocardial infarcts^18^), different tissue-type interfaces (e.g., bone-ligament^19,20^), and tumor-stroma interfaces.^21,22^ Whereas mechanical manipulation after tissue formation enables studies of how established tissues adapt to changing strain in exercise adaptation^23,24^ and disease progression.^25^ Several approaches have been developed for creating spatially patterned tissues, each with distinct capabilities and constraints. 3D bioprinting methods,^26–29^ such as extrusion^30^ and laser-assisted techniques,^31^ can produce complex geometries but typically require material-specific optimization: shear-thinning properties for extrusion, precise viscosities for inkjet deposition, or photopolymerizable components for light-based printing.^32^ Recent advances such as FRESH bioprinting have expanded the range of printable materials,^33,34^ while various tissue engineering platforms have successfully incorporated mechanical stimulation.^35^ Open microfluidic approaches using capillary-driven flow offer an alternative that bypasses material specific requirements while also allowing for the integration of mechanical stimulation.^14,36–45^

Exemplifying this approach, we previously developed Suspended Tissue Open Microfluidic Patterning (STOMP), a platform that allows for spatial patterning of suspended tissues through capillary pinning features in open microfluidic channels^44^ and enables multiregional constructs to study tissue interfaces. However, STOMP’s channel geometry was designed for string-like tissues suspended between two posts; this new approach allows for larger and more complex architectures and tissues. These geometries include larger tissue “patches”, which has implications in regenerative medicine applications (e.g., skin^46,47^ or cardiac patch^48–50^) and nonplanar suspended geometries expanding tissue modeling to curved surfaces, potentially relevant for modelling tissues such as gut epithelium, cervix, and cornea.

Here, we introduce Suspended Tissue Engineering with Assemblable Microfluidics (STEAM), a modular platform that integrates spatial patterning, mechanical manipulation, and portability through removable fluidic channel assemblies and portable tissue hook device systems. STEAM employs stackable 3D printed components, including a base patterning rail, tissue hook device that also serves as the channel walls, and top patterning rail, that temporarily form a fluidic channel for tissue fabrication via spontaneous capillary flow (SCF)^51^ of ECM-precursors. After gelling, the stacked assembly can be disassembled, leaving tissues suspended between portable tissue hooks, which serve as both attachment points and enable mechanical actuation. This separable fluidic channel design assists patterning rail removal by eliminating the requirement for tissue compaction, a feature utilized in the STOMP system to remove the patterning device from some tissue types. Further, the tissues remain suspended and attached to the tissue hook device(s); a user can pick up the tissue hook device to move the suspended constructs from patterning to culture to microscopy as needed without disrupting the tissue. Since tissue constructs can be moved around in this manner, they can also be stacked on top of each other, thus generating multiregional, and multilayered constructs.^43,44^ Additionally, STEAM is relatively material agnostic; unlike in many 3D bioprinting or hydrogel patterning methods that require specialized materials, STEAM uses SCF to pattern suspended tissue. Therefore, any material that transitions from a liquid precursor to a gel state (in a timespan long enough to flow into the channel), is theoretically possible to use with STEAM. The STEAM platform enables: (1) 3D geometries including planar structures in a rectangular shaped patch, and complex nonplanar configurations through customizable channel shapes, (2) multiregional patterning via sequential pipetting with capillary pinning features controlling fluid boundaries,^44^ and (3) up to 50% uniaxial static strain application by repositioning tissue hook devices on an incremental stretching device. As a proof-of-concept, we use C2C12 mouse myoblasts embedded in fibrin in the STEAM system and apply up to 50% strain, resulting in strain-induced myotube alignment. By integrating capabilities including spatial patterning, post-fabrication mechanical manipulation, and tissue portability, STEAM provides a unified, customizable suite for investigating how spatial organization, tissue architecture, and mechanical forces regulate tissue behavior in mechanically responsive, spatially heterogeneous tissues.

## RESULTS

### STEAM achieves geometric and mechanical control over suspended tissue architecture

STEAM generates suspended tissues and provides control at the intersection of tissue mechanics and 3D cell patterning. STEAM utilizes a stackable microfluidic assembly where a liquid ECM-precursor with embedded cells is pipetted into the assembled channel (Figure 1a, dimensions in Figure S1). Flow of the precursor in the channel occurs via surface tension-driven forces, and enables tissue patterning. The shape of the suspended tissue is dictated by the shape of the top patterning rail and the base patterning rail of the microfluidic channel and, when assembled together, the tissue hook device completes the channel (Figure 1b). After the ECM-precursor has gelled, the microfluidic channel can be disassembled, leaving a suspended tissue between the tissue “hook” anchor points (Figure 1c).

**Figure 1:**
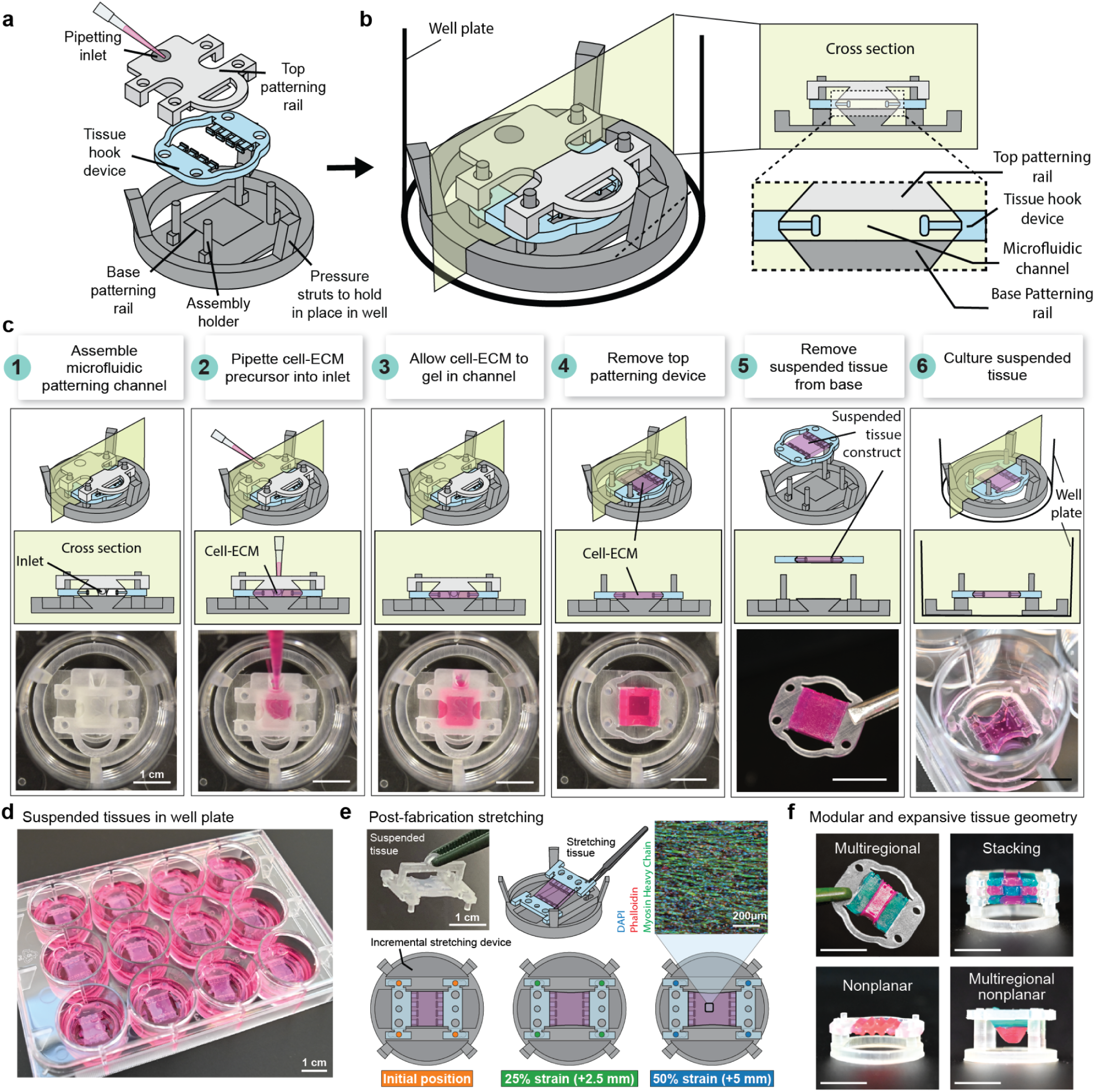
Suspended Tissue Engineering with Assemblable Microfluidics. **a** Exploded view of microfluidic patterning assembly, with the base patterning rail on the bottom, the tissue hook device stacked next, and the top patterning rail on top. **b** Full patterning assembly (left), with inset (right) showing a cross section of the fluidic patterning channel. **c** Schematic of patterning processes (top), with corresponding photos patterning agarose colored with india ink dye (pink). Bottom right most image is of a planar patch made with fibrin gel in a suspended holder within a 12-well plate. All scale bars are 1 cm. **d** Full 12-well plate of suspended planar patches made with collagen generated with STEAM. Scale bar is 1 cm. **e** Demonstration of post tissue fabrication method for applying strain to STEAM generated tissues. Top right image is a maximum projection confocal image of C2C12 mouse myoblasts differentiated into myotubes and stained for myosin heavy chain (green), phalloidin (red), and DAPI (blue). Scale bar is 200 µm. **f** Array of modular patterning abilities with expanded geometries including planar and nonplanar constructs, multiregional patterning of planar and nonplanar constructs, and stacking of separately patterned constructs. All scale bars are 1 cm.

The STEAM patterning process is as follows: (1) first, a microfluidic patterning setup is assembled by stacking a tissue hook device on top of the base patterning rail, and then a top patterning rail on top of the tissue hook piece. The stacked devices align via four posts on the base patterning rail, labeled as assembly holder in Figure 1a. The stacked assembly, made up of the base patterning rail, tissue hook device, and top patterning rail, comprise the “floor”, “walls” and “ceiling”, respectively, of a fluidic channel for hydrogel patterning. (2) Next, cell-laden ECM-precursor is pipetted with a manual pipette into the inlet attached to the top patterning rail to generate the suspended tissue construct. The inlet helps guide flow into the channel, but flow of the liquid ECM-precursor within the channel is governed by SCF.^51,52^ (3) After the patterning process is complete, the assembly is put in the incubator for cell-laden ECM-precursor gelation. (4) Once tissue gelation is complete, the temporary fluidic patterning assembly is disassembled. First, the top patterning rail is removed, then the tissue is lifted via the tissue hook device from the base patterning rail and transferred to a separate holder which allows for a suspended tissue culture in well plates (Figure 1d). Depending on the tissue type, patterning rail surfaces can be surface treated to aid in the removal process. The full tissue patterning process is illustrated in Figure 1c, and movies showing the patterning (Movie S1) and removal of the patterning rails (Movie S2) with a planar patch are available in the Supplementary Information. Critically, in addition to allowing the formed tissue to be placed in a suspended tissue culture, the portability of the generated tissue afforded by the tissue hook device enables moving living tissues between experimental platforms while preserving their architecture.

The suspended tissue architecture also enables several unique experimental capabilities after initial patterning. STEAM adds the ability to mechanically manipulate tissues after fabrication; the tissue hook system can be integrated with mechanical stimulation to apply controlled stretch to suspended tissues via an incremental stretching device and separated tissue hook devices (see static stretching of STEAM-generated tissues section below for more details), allowing for *in-vitro* strain-loading conditions that are important for mechanosensitive tissues like muscle, tendon, or cardiac tissue (Figure 1e).

The assemblable patterning method enables versatile geometric control through modular components that can be adapted for specific applications. One such adaptation is the implementation of pinning features in the patterning rails which enable multiple regions within a single construct, further increasing the dimensionality of patterning conditions. Furthermore, modifying the geometry of the top and base patterning rails enables control over the *z*-dimension, allowing the creation of any tissue construct from a planar tissue “patch” to more complex nonplanar geometries with or without multiregionality (Figure 1f). The modularity STEAM offers enables sophisticated tissue assembly strategies, such as culturing different cell populations under distinct conditions separately, then stacking them to create multilayered constructs with defined tissue-tissue interfaces. Additionally, by combining multiregional patterning with stacking capabilities (Figure 1f), STEAM enables the design of complex 3D cell signaling experiments with a temporal component, where different layers can be added or removed at different timepoints. Overall, this combination of geometric patterning, modular handling, and controlled mechanical stretching capabilities positions STEAM as a versatile platform for engineering complex tissue models.

### The addition of pinning features allows for multiregional control in STEAM tissues

SCF describes the surface tension driven transport of fluids through various open, semi-open, and closed microfluidic channels without requiring external pressure sources or actuators.^51–53^ In STEAM’s temporary channels, the driving mechanism of the fluid flow arises from the advancing meniscus of the hydrogel precursor solution, where capillary pressure generated by the curvature of the liquid interface pulls the fluid forward through the channel. This spontaneous flow only occurs when the distance between the top and bottom patterning rails (e.g., the height of the fluidic channel) is sufficiently small that a negative capillary pressure builds at the meniscus. In the context of STEAM, SCF provides a simple mechanism for patterning hydrogel precursors into defined 3D tissue constructs. By leveraging surface tension forces, pre-gel liquid solutions can be delivered precisely through assembled channels formed between patterning rails using only manual pipettes. This eliminates the need for complex pumping systems while maintaining the spatial control necessary for engineering relevant tissue architectures.

To achieve spatial control over the SCF driven fluid flow of precursor hydrogels in STEAM, we incorporated pinning features adapted from our established microfluidic patterning methods.^43,44^ Specifically, we implemented similar pinning feature geometries that were characterized in our previous STOMP platform to cessate the flow of hydrogel precursor in our assembled channel to enable multiregional patterning within STEAM tissues.^44^ The geometry comparison between STEAM and STOMP is illustrated in Figure 2a and 2b, respectively. Briefly, in the STOMP system the pinning features are perpendicular to the direction of flow (Figure 2b), and the advancing fluid front stops at the pinning feature. Unlike in the STOMP system, the advancing fluid front moves parallel to the pinning features in STEAM, and acts as a flow guiding feature (Figure 2a). The pinning feature creates a virtual channel “wall” preventing flow into the adjacent region (Movie S3). We use similar flow guiding features in our previous work on layer-by-layer hydrogel patterning with open microfluidic systems.^43^

**Figure 2:**
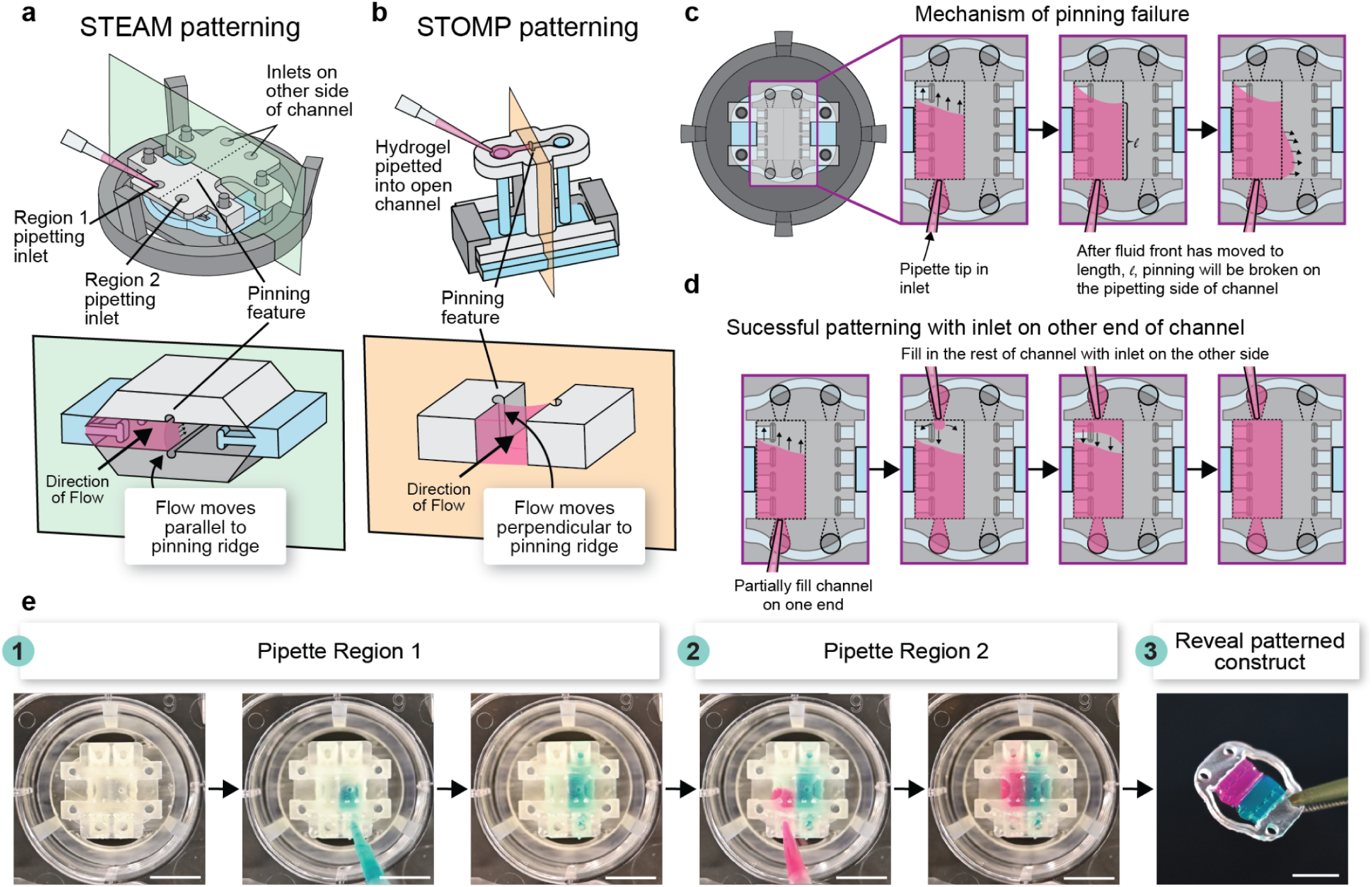
Multiregion patterning in STEAM. **a** STEAM patterning assembly (top) with an isometric cross-sectional view (bottom) showing flow moving parallel to the pinning ridge, compared to the configuration of the **b** STOMP patterning assembly (top) with an isometric cross-sectional view (bottom) showing flow moving perpendicular to and halting at the pinning ridge. **c** Illustration of the observed mechanism of pinning failure, where flow moves down the channel and pins along the pinning ridge until a given length, *ℓ,* where the fluid front then breaks and flows past the pinning ridge on the pipetting side of the channel. **d** The utilization of a second pipetting inlet on the opposite end of the channel mitigates this pinning issue and allows for successful patterning throughout the length of the channel. **e** Top down workflow of patterning two regions in STEAM with colored agarose. All scale bars are 1 cm.

A critical consideration when designing pinning features for STEAM is understanding the conditions under which they fail, as depinning events can compromise spatial control over the patterned tissue. In STEAM, it is theorized that a depinning event occurs after the fluid front reaches a particular length, ℓ, along the channel, effectively losing spatial control of the tissue (Figure 2c). The depinning happens near the inlet of the channel and likely occurs due to an increase in pressure at the inlet. To mitigate this, a second inlet mirrored on the other side of the patterning rail can be used to fully fill each region with the appropriate amount of ECM-precursor (Figure 2d). Since the STEAM system has a semi-open configuration, it is possible to add inlets in multiple locations along the channel to reduce the travel distance of the fluid and allow successful pinning to occur, therefore increasing the number of possible pinning geometries. The length the fluid front can reach before de-pinning (ℓ) was observed to be different based on the fluid identity. Therefore, this multiple inlet approach may only be needed in specific scenarios based on channel geometry and desired ECM properties; this was also a key consideration in the STOMP system.^44^ The two previously characterized pinning feature designs in the context of STOMP include the concave “cavity” pins and the convex “vampire” pins, which we have now engineered on the top and base patterning rails in STEAM.^44^ The example workflow in Figure 2e utilizes “cavity” pinning features; both pinning geometries are depicted in Figure S2. An example of the removal process for a two region planar patch is included in Movie S4.

### Modular stacking of STEAM patches enables spatial diffusion dynamics

In addition to spatial patterning via capillary pinning, STEAM patches can be stacked and separated at will to create modular, spatially and temporally defined multi-culture systems. This enables both vertical gradients between stacked layers and horizontal gradients within layers, critical for modeling native tissue environments where distance from signal sources drives differential cellular responses. By tuning the distances between stacked tissue sections, more complex signaling gradients can be generated than are achievable through spatial patterning alone. The modular nature of STEAM further allows for easy separation and analysis of individual layers, and patches can conceivably be added or removed at defined timepoints to interrogate dynamic signaling processes. In previous work, we developed a similar modular tissue stacking technique and demonstrated cellular signalling between layers of tissue stacked together.^37^ STEAM builds upon this by enabling spatial patterning *within* a hydrogel patch, and further expands the geometry available for stackable structures.

To demonstrate tunable gradient generation, 10 kDa fluorescent Texas Red dextran was chosen due to its size being comparable with other researched cellular signaling proteins such as the chemokine ligand CXCL10^54^ and thioredoxin.^55^ For this experiment, the dextran was patterned into either a single-region patch (Scheme 1) or one region of a two-region collagen patch, with the adjacent region left as blank collagen (Scheme 2). A second blank collagen patch was stacked on top of both configurations (Figure 3a). STEAM-generated collagen patches were patterned separately, stacked together, incubated to allow diffusion between and within patches, and then separated for analysis (Figure 3b-c). Confocal fluorescent imaging and analysis of both the top and bottom STEAM-generated patches before and after the stacking process revealed diffusion of dextran molecules between and within stacked STEAM patches (Figure 3d). For the single region patch configuration (Scheme 1), we observed dextran diffusing across the X-Y axis of the total construct from the initial patterned patch to the stacked blank patch (Figure 3di). For the two region initial patterning configuration (Scheme 2), a spatial gradient of dextran molecules was observed in both patches, the initially patterned patch and the initially blank patch (Figure 3dii). A workflow for how dextran diffusion was quantified from confocal fluorescent images and a second replicate for each initial configuration is shown in Figure S3 and Figure S4, respectively.

**Figure 3:**
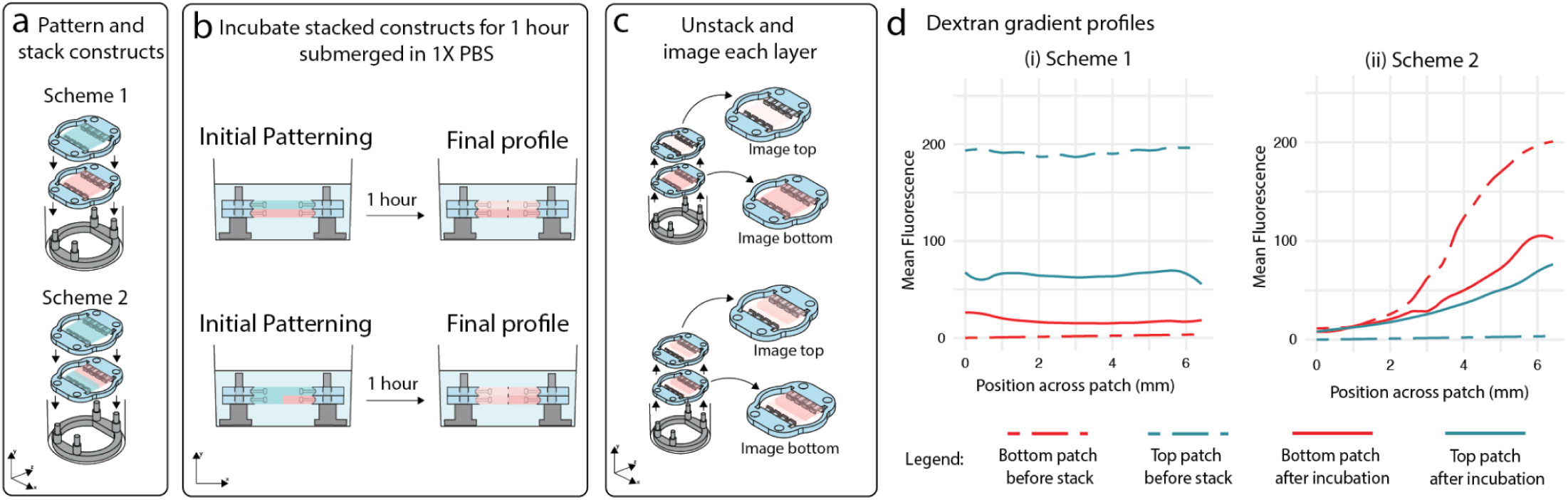
Experimental design for spatial diffusion of dextran via STEAM. **a** STEAM planar patch constructs were patterned and stacked on top of each other in a 12 well plate. Experimental scheme 1 was a dextran-infused single region collagen patch with a blank collagen patch on top while experimental scheme 2 was a two region patch with dextran-infused collagen on one side and blank collagen on the other with a blank collagen patch on top. **b** Stacked constructs are submerged in 1X PBS for 1 h to allow for diffusion of dextran. **c** The constructs were then unstacked and confocal imaged separately. **d** Fluorescent profiles were generated from intensity of dextran of the collagen patches before incubation (dashed line) and after incubation (solid line), experimental scheme 1 revealed diffusion with an increase of dextran in the top collagen patch (blue) and a decrease in the bottom collagen patch (red). For experimental scheme 2, the same trend as scheme 1 was revealed with an increase of dextran in the top collagen patch (red), however, a spatial gradient was detected due to the initial two region patterning of the bottom collagen patch (blue) and initial placement of dextran.

### Static stretching can be applied to STEAM-generated tissues with a simple mechanism

Beyond spatial patterning and modular stacking, STEAM incorporates mechanical manipulation capabilities critical for engineering mechanosensitive tissues. Many tissues require mechanical stimulation for proper development and function: skeletal muscle cells require uniaxial tension for myotube alignment and maturation,^56,57^ tendons develop under tension,^58^ and vascular smooth muscle cells remodel in response to stretch.^59,60^ STEAM adds stretching capability through a modified tissue hook system that enables mechanical strain application after tissue patterning. In this configuration, the tissue hook devices are designed as two independent components rather than a continuous frame, with each set of hooks containing pegs on their bottom surface, allowing for the final patterned tissue to be held at a particular amount of strain when interfaced with the corresponding bottom hole of the incremental stretcher device (Figure 4). This disconnected design serves dual purposes: it maintains tissue suspension during culture while enabling lateral displacement for fixed, static strain application.

**Figure 4:**
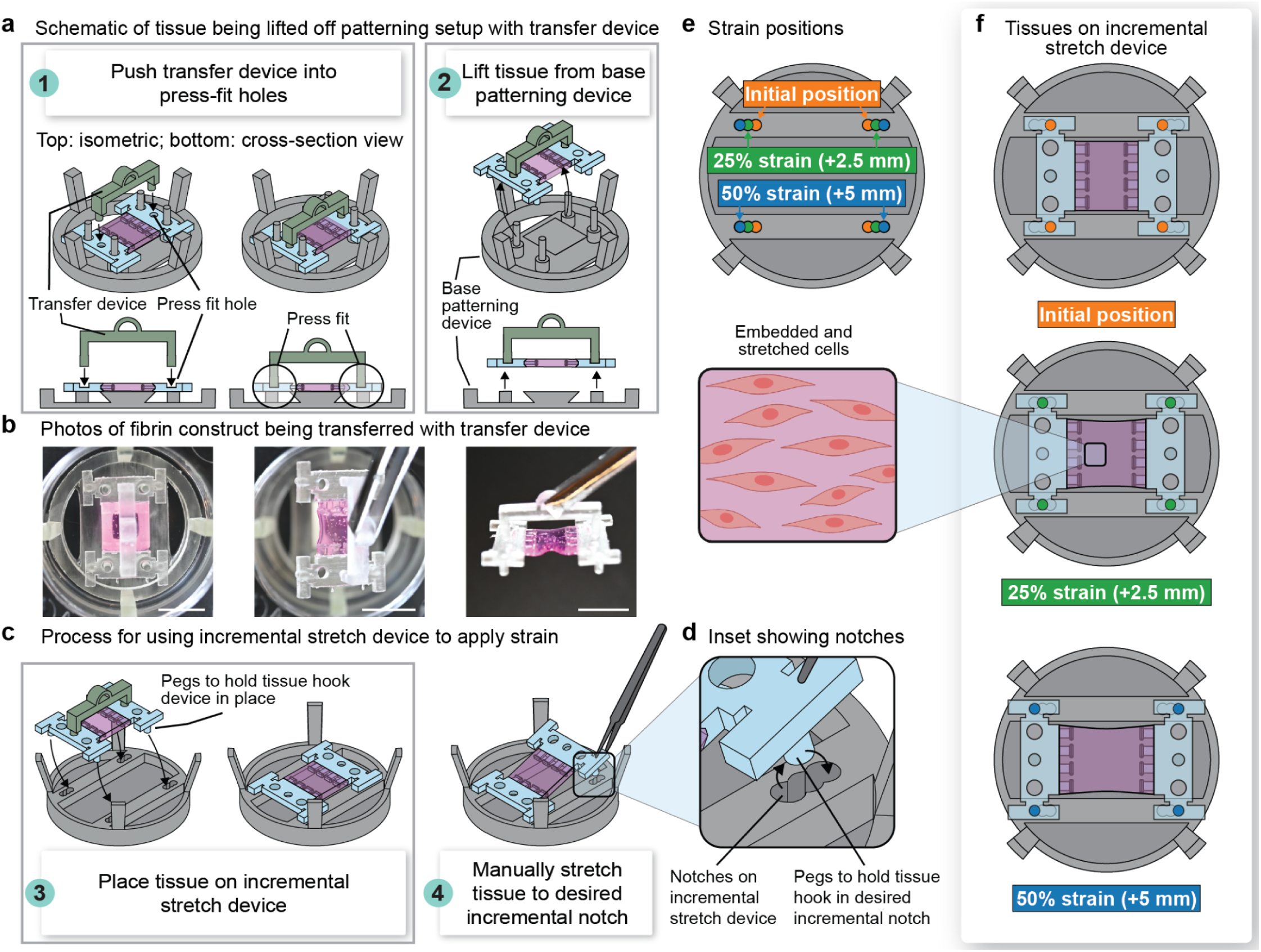
Static stretching via STEAM. **a** Schematic workflow for lifting tissue off the patterning setup using the transfer device. The transfer device uses a press-fit feature to temporarily attach to and stabilize the tissue hooks for transfer. Top schematics show an isometric view, and bottom shows a cross-sectional view. **b** Photos of the transfer device being used to lift a fibrin planar construct from the patterning device. All scale bars are 1 cm. **c** Schematic workflow of placing the tissue on the incremental stretch device by aligning the pegs on the bottom of the tissue hook devices with the “initial position” notches on the incremental stretch device. Next, the tissues can be manually stretched by lifting the tissue hook device and moving it so the pegs are placed in a different notch. **d** Inset showing notches on the incremental stretch device. **e** Top down schematic showing the notch positions of the incremental stretch device for the initial position (0% strain), 25% strain condition, and full 50% strain condition. **f** Top down schematics showing a planar tissue construct stretched at each strain condition (0%, 25%, and 50%) with an inset graphic of embedded and stretched cells.

This mechanical loading leverages STEAM’s modular design principles. Following tissue patterning and gelation, a transfer device with press-fit features engages the top of each tissue hook, temporarily connecting and stabilizing them during transfer due to the slight size difference in the diameter of the transfer device and the associated press-fit feature on the tissue hook devices (Figure 4a-b). The tissue is then moved to a static incremental stretching device, where the bottom pegs secure each hook device in a predefined position (Figure 4c-d). Once transferred, controlled strain is applied by sequentially repositioning the tissue hooks using tweezers into holes on the incremental stretching device spaced at predetermined distances (Figure 4e-f). This is done one tissue hook at a time, first by lifting one peg on the bottom and inserting it into the next hole of the incremental stretching device before moving on to the other peg. This incremental approach prevents tissue tearing while achieving a stretch up to a 5 mm increase (50% strain for the 10 mm tissue patches demonstrated in this manuscript). We refer to this configuration as “static” strain, as it will hold the tissue in the stretched configuration continuously rather than cycling between different strain conditions. However, tissues can be manually moved from one strain level to another incrementally as experimentally needed in the current configuration, and the tissue hook devices are also amenable to being attached to a motor and cyclically strained in future iterations.

### STEAM static stretching enables increased alignment of engineered skeletal muscle tissue patches

Muscle tissues *in vivo* exhibit a high degree of alignment, enhancing mechanical and cellular function.^61,62^ Proper cellular alignment in mature skeletal muscle tissue is crucial for efficient contraction and force generation. During myotube formation, a critical step in skeletal muscle maturation, skeletal muscle myoblast cells elongate and fuse to form multinucleated myotubes.^63,64^ *In vitro*, organized alignment among myotubes more accurately recapitulates native tissue architecture and function, and is a critical indicator of muscle maturity.^63,64^ When modeling muscle tissues *in vitro*, mechanical stretching of the ECM has been shown to improve tissue remodeling and differentiation in skeletal muscle cells by promoting increased muscle maturation. This is often measured by the myotube fusion index, increased myonucleation, uniaxial alignment of myotubes, and increased functional force output.^24,25,61,62,65–68^ To demonstrate the functional impact of the mechanical manipulation achievable in STEAM, we applied static strain to C2C12 mouse myoblasts fully embedded in suspended fibrin ECM. When using STEAM, the entire patch, including the ECM, is put under a set strain by stretching the tissue uniaxially to increase the final length.

We hypothesized that applying an increased strain (25% and 50%) to the tissue using the STEAM static stretch system would induce more alignment in our engineered muscle tissues compared to an unstretched STEAM suspended tissue patch left at the initial position (Figure 5a). To visualize cellular structures, tissues after 5 days of differentiation and post-fabrication strain were immunostained with myosin heavy chain antibody for myotube formation, phalloidin for F-actin in the cellular cytoskeleton, and DAPI for cellular nuclei (Figure 5b). Nuclear alignment, or the orientation of the long axis of the nuclei (where nuclei are fit to an ellipsoid shape),^68^ is more distributed towards the direction of stretch (defined as 0°) in the higher strain tissues (25% and 50%), suggesting strain-induced alignment (Figure 5c). We found that both the 25% and 50% conditions were significantly different from the control condition (*p* ≤ 0.01) when the mean nuclear alignment angle was compared, however the 25% and 50% conditions did not differ significantly from each other (*p* > 0.05) (Figure S5). A conceptual illustration of the angle measured in the nuclear alignment analysis is also provided in Figure S5. Utilizing two-dimensional fast Fourier transform (2D FFT) analysis of the confocal fluorescent images (see Figure S6 for more detail), we saw an increase in myotube alignment from 0% to 25% strain and from 25% to 50% strain (Figure 5d). The 2D FFT analysis includes a radial sum function to create a plot of the representative principal angles of myotubes in a 360° space corresponding to the image.^65^ Therefore, the level of myotube alignment is determined by comparing the height and width of the peaks obtained, with a singular peak corresponding to a high degree of myotube alignment along an angle in the radial summation alignment plot. The level of myotube alignment is further characterized by comparing the height and width of the peaks obtained, with a narrower width and sharper peak shape corresponding to more aligned tissues; in our system, this peak would be at 90° (Figure 5d).^65^ Unlike in the nuclear alignment analysis where the direction of stretch is defined as 0°, the direction of stretch in the 2D FFT alignment plot is defined as 90°. Notably, we see singular peaks across the various stretch conditions which is expected as even the control (0% strain) is grown in suspension and therefore experiences an inherent degree of tension; however, as strain is applied, we obtain narrower peaks, indicating a higher degree alignment (Figure 5d). We found that adding strain to the suspended STEAM generated tissue patch improved myotube alignment, validating that STEAM’s strain application maintains biological functionality while providing the mechanical cues necessary for tissue development.

**Figure 5:**
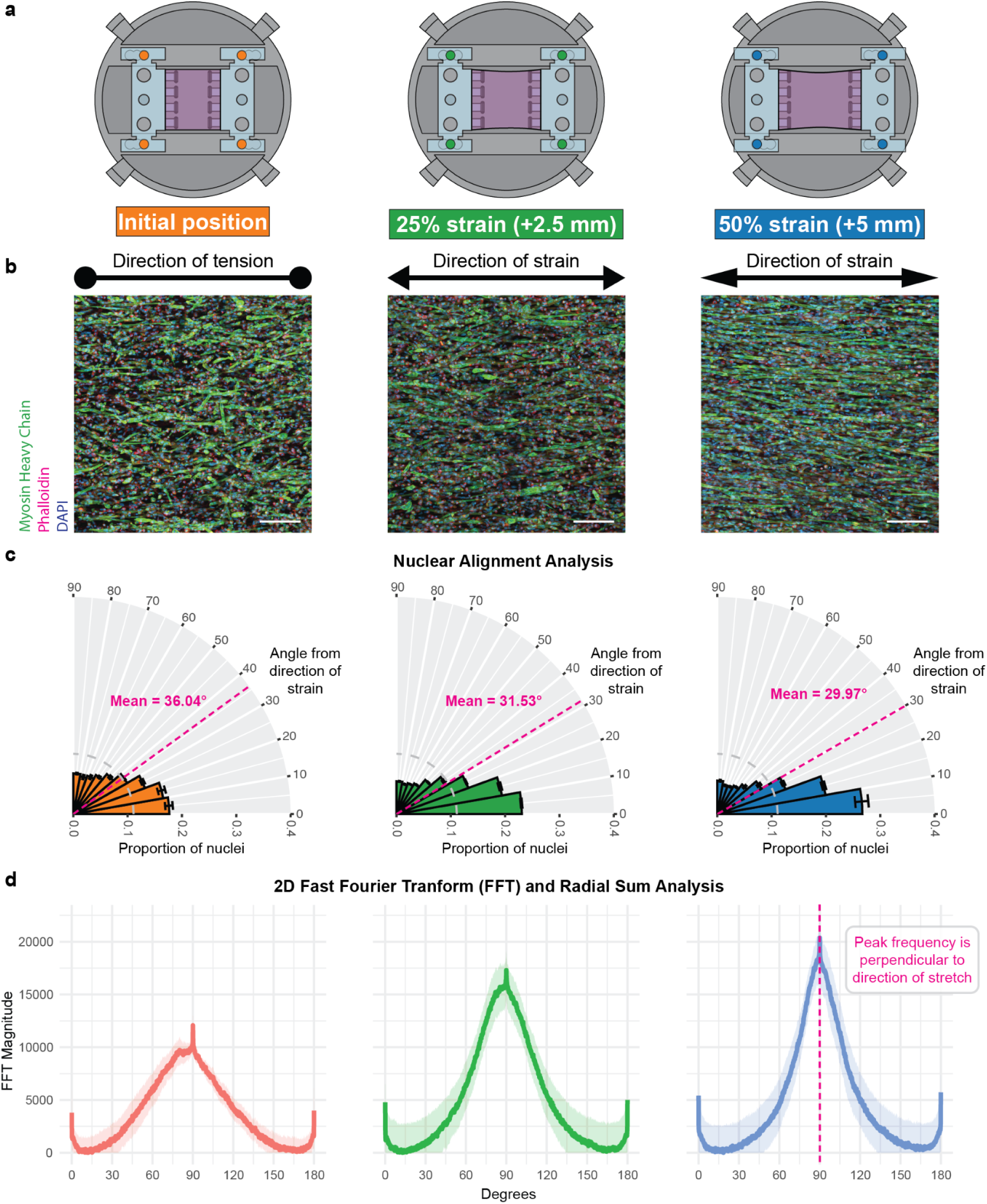
STEAM muscle tissue patch shows increased alignment with applied strain. **a** Schematic showing the three positions of strain applied, including 0% (initial position, left), 25% (middle) and 50% (right). **b** Representative confocal images of tissues after 5 days of differentiation and post-fabrication applied strain (Left 0%, middle 25%, right 50%). Confocal images show a maximum projection of C2C12 cells stained for the nucleus (DAPI, blue), myotube formation (myosin heavy chain, green), and the cytoskeleton (phalloidin, red) for each of the conditions. Scale bar is 100 µm. **c** Nuclear alignment analysis of the three strain conditions (Left 0%, middle 25%, right 50%). Error bars represent the standard deviation of three separate tissues from a single experiment. 0° is defined as the direction of stretch. Illustration of nuclear alignment angle measurement is found in Figure S4. **d** 2D Fast Fourier transform (2D FFT) and radial sum analysis of the three strain conditions (Left 0%, middle 25%, right 50%), with shading representing standard deviation of three separate tissues from a single experiment. 90° is defined as the direction of stretch as peak frequency is calculated perpendicular to the direction of stretch in a more aligned tissue.

To investigate the impact of strain on muscle differentiation and maturity we also assessed mRNA expression levels. Muscle differentiation markers *MyHC4*, *MYOG*, *DES*, and *CKM* were compared to undifferentiated 3D engineered muscle tissues indicating successful differentiation of muscle cells when cultured with these devices (Figure S7, Table S1,^40^ and Supplementary Text). When mRNA levels are compared within differentiated control (no added strain) and varied strain, there is no marked difference between conditions (Figure S8). Myotube diameter was also assessed as a measure of myotube maturity across the strain conditions, which showed no significant difference (see Figure S9, and Supplementary Text). Overall, degrees of strain did not affect muscle cell differentiation and myotube formation; all suspended tissues in the STEAM system (regardless of added strain) expressed *MyHC4*, *MYOG*, *DES*, and *CKM* via mRNA expression and myosin heavy chain and phalloidin via immunostaining.

Collectively, these morphologies demonstrate that static mechanical strain applied through the STEAM platform effectively drives skeletal muscle tissue alignment in suspended engineered muscle tissue patches. The response to strain indicates that STEAM’s mechanical manipulation preserves and enhances the mechanosensitive responses essential for muscle development. Importantly, the achievement of 50% strain without tissue failure validates the robustness of the suspended tissue architecture, while the progressive myotube alignment improvements from 25% to 50% strain indicate that the system operates within the dynamic range of cellular response to mechanical stimulation. These results establish STEAM’s capability to provide controlled mechanical stimulation to fully embedded 3D muscle tissues, distinguishing it from conventional 2D substrate stretching approaches such as where cells are cultured on top of an ECM or other stretchable substrate (e.g., polydimethylsiloxane (PDMS) or silicone).^69,70^

### STEAM enables expanded geometric control through nonplanar architectures and modular stacking

Beyond planar patches, STEAM generates 3D tissue architectures through modification of the shape of the top and bottom patterning rails. When these rails incorporate curved surfaces, the resulting tissues adopt nonplanar configurations while maintaining suspension between tissue hooks. The cross sections of a planar patch and nonplanar wave geometry patterning setup are illustrated in Figure 6a. The wave geometry shown in Figure 6a-b shows how coordinated shaping of patterning devices produces suspended tissues with controlled curvature, expanding the suspended tissue geometric repertoire beyond planar patches and string-like architectures.

**Figure 6:**
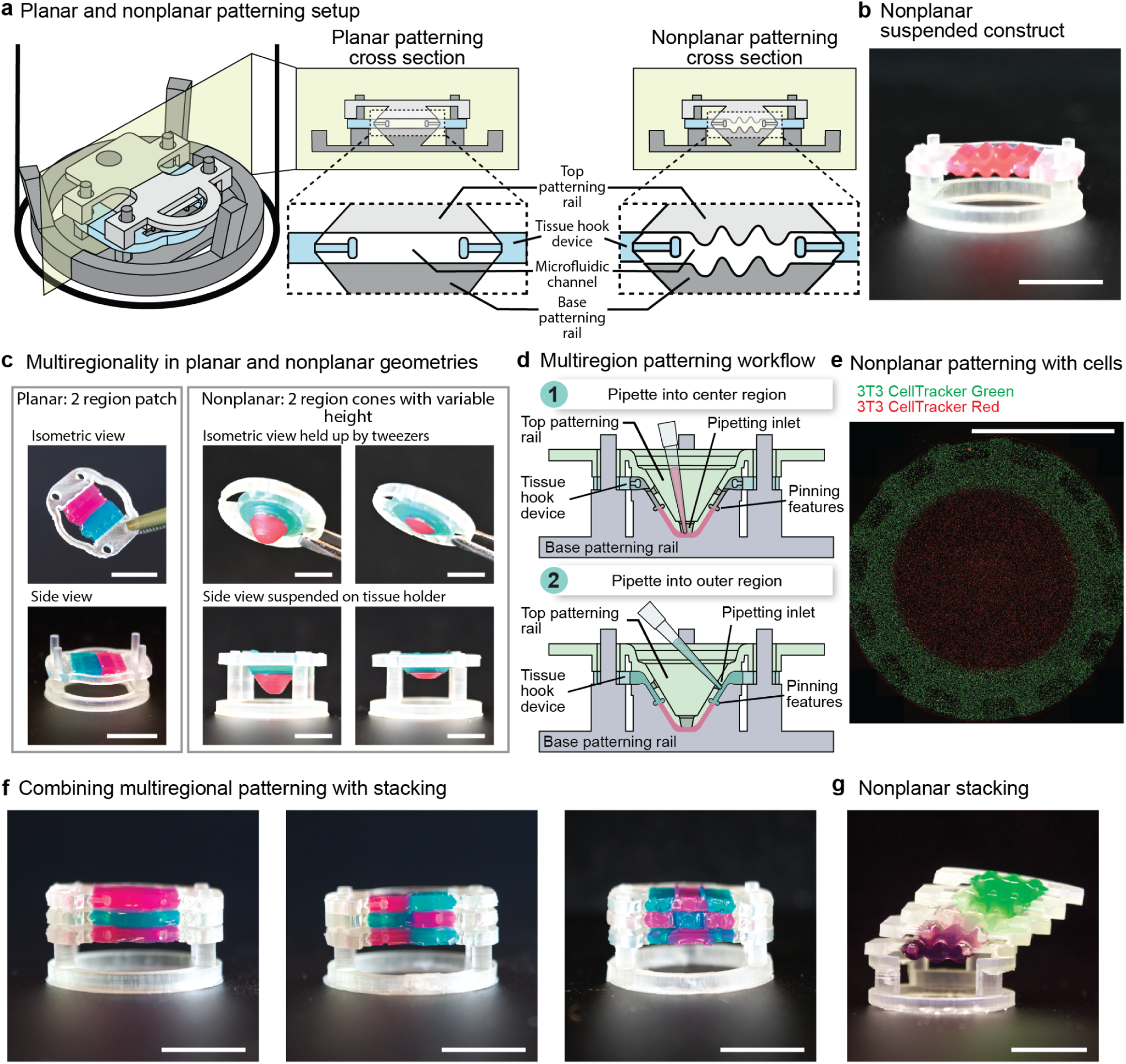
Expanded patterning geometries with STEAM. **a** Patterning setup with STEAM showing the cross section for a planar (left) patch geometry and a nonplanar (right) wave geometry. **b** Nonplanar wave geometry suspended construct made from colored agarose. Scale bar is 1 cm. **c** Multiregional patterning in planar and nonplanar geometries with colored agarose. Constructs in a planar geometry (left) and a nonplanar geometry, specifically a two region cone-shape with varying heights (right). All scale bars are 1 cm. **d** Schematic showing process for patterning a two region cone. **e** Representative confocal image (maximum projection) showing successful patterning of a two region cone geometry with mouse 3T3 fibroblast cells dyed with CellTracker red (inner region) and CellTracker green (outer region). Scale bar is 1 cm. **f** Multiregional planar constructs combined with stackability. Examples of a single region construct (left), two region construct (middle), and a three region construct (right). All scale bars are 1 cm. **g** Nonplanar stacking of colored agarose wave geometries with diffusion between layers. Initially, the top-most layered structure was dyed green, the bottom-most layered structure was dyed purple, and the middle three layers were clear. Image taken 2 hours after stacking. Time lapse of diffusion included in Movie S5. Scale bar is 1 cm.

Multiregional patterning extends to nonplanar architectures with incorporation of pinning features used in planar structures to create two distinct regions of cone-shaped constructs (Figure 6c-d). The cone geometry maintains spatial segregation of differentially labeled 3T3 mouse fibroblast cells (CellTracker red in inner cone region, CellTracker green in outer cone region) (Figure 6e) while introducing variation in *z*-height from center to periphery. This capability allows modeling of cellular interfaces that exist across curved surfaces, such as those present in tubular organs or at tissue boundaries with complex topographies.

STEAM’s modular architecture facilitates assembly of multilayer 3D suspended tissue constructs through vertical stacking of patches. Figure 6f demonstrates progressive complexity achieved by stacking constructs containing one, two, or three patterned regions. Since each layer is patterned independently before assembly, different cell populations can be cultured under their optimal conditions prior to combining. The tissue hook device maintains each layer’s geometry during stacking while preserving the ability to separate layers for individual analysis. This modularity enables temporal control over tissue assembly, where layers can be added or removed at specific timepoints to study dynamic cell-cell interactions or sequential tissue formation processes.

The combination of nonplanar geometries with vertical stacking produces 3D gradients that follow the construct architecture. In Figure 6g, five wave shaped agarose constructs were assembled with colored food dye in the top (green) layer and bottom (purple) layer, with three clear layers stacked in between. After 2 hours, diffusion created a gradient of colored dye both vertically through the stack and laterally along the wave contours, demonstrating how STEAM can generate spatially complex diffusion patterns (Movie S5). Such controlled nonplanar gradients could be used to study how signaling molecules distribute through curved tissues or how tissue geometry influences molecular transport.

## DISCUSSION

Suspended tissue models have long been shown to recapitulate tissue scale mechanics, but achieving spatial heterogeneity within mechanically functional systems remains challenging. STEAM enables experimental control over suspended tissues by combining spatial patterning and mechanical manipulation. The assemblable fluidic channel design allows complex spatial patterns to be created via SCF, and then the channel components are lifted away, leaving tissues suspended on hooks that serve as both anchors and mechanical actuators. This approach transforms suspended tissues from static constructs into dynamic experimental systems that can be spatially controlled, mechanically stretched, transferred between platforms, and assembled into multilayer structures. As a proof-of-concept, our muscle tissue experiments demonstrate how modulating mechanical conditions post-fabrication can provide additional mechanical cues to drive cellular organization. Progressive myotube alignment with increasing strain validates STEAM’s capability to model how established tissues adapt to changing mechanical loads. By combining temporal control over mechanics with spatial patterning capabilities, STEAM addresses a historical challenge in tissue engineering.

STEAM builds upon our group’s recent advances in surface tension driven tissue patterning while expanding the geometric possibilities. Similar to our recently developed STOMP platform,^44^ STEAM harnesses SCF to pattern ECM-precursors in a relatively material agnostic manner, eliminating the need for specialized pumping equipment or material specific protocols. However, STEAM extends the geometric repertoire beyond STOMP’s string-like configurations to encompass planar patches and complex nonplanar architectures. Further, STOMP generally works best with tissues that are able to compact away from the channel walls, allowing for simple removal of the patterning device. STOMP can work with noncompactible cell-ECM combinations, but this requires a device with additional patterning steps and degradable channel wall. STEAM’s removable channel architecture does not rely on tissue contraction and therefore eliminates this limitation.

While STEAM demonstrates versatile tissue patterning capabilities, several opportunities exist for system enhancement. For example, the current mechanism for stretching tissues relies on manual manipulation for strain application, and tissues are held at a static strain. This precludes taking functional measurements, such as those that can be achieved in a two post suspended tissue system. In these systems, tissues are suspended between two posts made from a flexible material such as PDMS. This allows for the tissue to move the post upon contraction, thus allowing for functional measurements by calculating the distance the post was deflected. However, there are additional design approaches that could be integrated to STEAM’s tissue hook system, such as integrating spring-like structures, to allow for similar deflection and functional force measurements. Integration with motorized actuators would enable automated cyclic mechanical stimulation, which is important for development of tissues that regularly undergo cyclic mechanical strain cues such as cardiac,^71^ bladder,^72^ and muscle tissue^57^. Since we did not incorporate cyclic stretching in our demonstration of muscle tissue patches, this may have contributed to the lack of significant differences in mRNA levels of certain differentiation markers and myotube diameter, as it has been demonstrated previously that cyclic stretching protocols achieve better maturation.^15,23,24,56,59,61,64^ Further, scaling considerations present both challenges and opportunities: flow via SCF restrains the height of each individual tissue, thus limiting the overall scale in the *z*-direction. Our stacking approach partially addresses this limitation, as layers of tissue can be generated and then stacked together to model a larger tissue. At larger scales, tissues require a perfusable “vascular” network to deliver oxygen and nutrients, which is the topic of ongoing work with the STEAM system.

The capabilities afforded by STEAM, including generating spatially heterogeneous tissue architectures that remain mechanically manipulable throughout their culture, distinguish STEAM as a platform that enables modularity and multifunctionality in suspended tissue engineering. The expansion to nonplanar geometries demonstrated in this work further extends STEAM’s experimental range, enabling investigation of tissues with curved architectures that better represent the complex 3D structures found *in vivo*. Combined with the modular stacking approach, these nonplanar capabilities allow creation of spatially complex tissue models. Furthermore, this modular approach would enable culturing of tissue layers separately before assembly, allowing for condition optimization for each tissue type before studying their interactions together. The reconfigurable stacking architecture also enables spatial interrogation of signaling dynamics between hydrogel layers. By stacking and subsequently separating STEAM patches, diffusion of fluorescent dextran molecules between discrete regions was quantified, demonstrating how future researchers could use STEAM to characterize the progression of molecular gradients across both vertical and lateral dimensions in a controlled, modular format. Furthermore the tissue hook system enables facile transfer of the tissue between various platforms. These could include transfer to microscope slides for precise interventions such as laser ablation of a specific region of tissue, controlled spatial crosslinking, or void patterning within intact tissues. Since the tissue remains suspended within the tissue hook device during transfer, its geometry can be preserved while gaining access to additional experimental platforms. Modular tissue engineering approaches have been explored,^73,74^ and multilayer assemblies have been achieved through various methods,^43,75^ however, STEAM’s suspended format provides a complementary approach with additional features of mechanical manipulation. By bridging the gap between simple suspended tissues and more complex engineered constructs, STEAM provides a foundation for studying how spatial organization and mechanical forces jointly regulate tissue behavior in mechanically active tissues.

## METHODS

### Device design, fabrication and sterilization

All STEAM patterning components were designed in SolidWorks 2025. Components were 3D printed out of clear V4 resin using Form 3/3B stereolithography 3D printers (Formlabs Inc.). Devices underwent a standard cleaning sequence to remove excess uncured resin. This sequence begins with agitation in isopropyl alcohol (IPA) for 20 and 10 min in two separate FormWash units (Formlabs Inc.). Devices were then transferred to beakers filled with fresh IPA for 30 min of sonication, after which devices were dried with pressurized air or set out on the bench until dry. Post-drying, devices are cured with ultraviolet light at 60 °C for 15 min in a FormCure (Formlabs Inc.). Prior to biological experiments with cells that involve stretching the tissues, the tissue hook pieces were treated by oxygen plasma using a Diener Zepto PC EX Type PB plasma treater (Diener Electronic, Germany). The plasma treatment helps keep the tissues attached to the tissue hooks during culture. Following plasma treatment, all device components are moved to a biosafety cabinet (BSC) for at least 15 min of UV to finalize the sterilization process After sterilization, the top and base patterning rails were incubated in a solution of 1% bovine serum albumin (BSA) for 1 h at room temperature as described previously^43,44^. After incubation, the 1% BSA solution was aspirated, and the patterning rails were allowed to fully dry prior to assembling.

### Patterning one-, two-and three-region STEAM patches with agarose

Patterning devices were assembled by placing the base patterning rail in a 6-well plate or on the bench, then the tissue hook device was placed onto the base posts, and finally the top patterning rail was placed on top. For patterning, low gelling temperature agarose (Sigma-Aldrich) was dissolved in deionized water to a concentration of 15 mg mL^-1^ and colored with india ink or food dye for visualization purposes (Dr. Ph. Martin’s or Spice Supreme, respectively). The agarose was heated to a liquid state (95 °C) before patterning with a standard pipette. For a single region patch, approximately 200 µL of agarose is pipetted into the inlet. For a two region patch, approximately 110 µL of agarose is pipetted into the first region and allowed to gel for 1-2 min before approximately 115 µL of agarose in another color is pipetted into the other inlet to form a contiguous two region construct. For a three region patch, approximately 65 µL of agarose is pipetted into both of the two inlets closest to the tissue hooks and allowed to gel for 1-2 min before approximately 65 µL of agarose is pipetted into the center inlet in another color to form a contiguous three region construct. All devices were then allowed to gel at room temperature (approximately 20-22 °C) for 5-10 min before patterning rail disassembly. All agarose structures used 1.5% wt/v gel. Images of the agarose constructs were then taken using a Nikon D5300 DSLR high resolution camera.

### Stacking and diffusion modeling with one-and two-region STEAM patches

The top and base patterning rails for both one-and two-region STEAM planar patch devices were incubated in a solution of 1% BSA for 1 h at room temperature. After incubation, the 1% BSA solution was aspirated and the patterning rails were allowed to fully dry prior to assembling. One-and two-region STEAM patterning devices were assembled by placing the base patterning rail in a 6-well plate, then the tissue hook device was placed onto the base posts, and finally the top patterning rail was placed on top. Texas Red dextran (10 kDa, Fisher) was chosen as the fluorescent marker for diffusion and was resuspended at 5 mg mL^-1^ in deionized water. One-and two-region STEAM patches were patterned with collagen. The collagen solution was prepared by mixing a stock of rat tail collagen type I (Corning) in 0.02 n acetic acid with deionized water, 10X PBS, 1M HEPES buffer solution, 7.5% sodium bicarbonate solution, and 1 N NaOH to achieve a final collagen density of 5 mg mL^-1^ and kept on ice to prevent gelation. Two liquid collagen solutions were prepared; one containing the dextran solution in place of the deionized water and one containing blank deionized water. For a single region patch, approximately 230 µL of collagen is pipetted into the inlet. This was replicated twice to create blank collagen STEAM planar patches for stacking and once to create a dextran-infused collagen STEAM planar patch. For a two region patch, approximately 65 µL of dextran-infused collagen was pipetted into a single inlet on each side of the first region and allowed to gel for 2-3 min before another 65 µL of blank collagen is pipetted into a single inlet on each side of the second region. These patches are incubated for 30 min at 37 °C for gelation. After gelation, confocal images were taken of each planar patch for baseline fluorescence. A single region dextran-infused collagen planar patch was then placed on a holder in a 12 well plate. A single region blank collagen planar patch was then placed on top and the whole construct was submerged in 3 mL of 1X PBS and incubated at 37 °C for 1 h. A two region planar patch where one region was dextran-infused collagen and the other region was blank collagen was also placed on a separate holder in a 12 well plate. A single region of blank collagen planar patch was then placed on top and the whole construct was submerged in 3 mL of 1X PBS and incubated at 37 °C for 1 h. After the 1 h incubation time, tissue stacks were disassembled and each planar patch was imaged via confocal microscope again to assess dextran diffusion within the stacked construct.

### Dextran diffusion quantification image analysis

The maximum projection confocal images of each patch were analyzed with an NIH ImageJ software to determine the spatial intensity of fluorescent dextran across the patch. To do this, five evenly spaced regions perpendicular to the direction of patterning were generated and the fluorescent intensity was averaged at each point across the length of the patch. The fluorescence profiles of each region were averaged and plotted to determine the average fluorescence profile of dextran across the length of the patch (see Figure S3). This method was chosen to denoise without changing the raw image data. One fluorescence profile per patch was then used to plot and visualize data in Figure 3. A second replicate for each initial configuration is shown in Figure S4.

### Cell culture maintenance - C2C12 mouse myoblasts

A mouse myoblast (C2C12) cell line was obtained from the American Type Culture Collection (ATCC). The cells were maintained in a tissue culture flask containing Dulbecco’s Modified Eagle Medium (DMEM) supplemented with 10% fetal bovine serum (FBS) and 1% penicillin-streptomycin at 37 °C, 5% CO_2_. Culture medium was changed every 48 h until cells reached around 40% confluency, whereupon the culture was rinsed with 1X phosphate buffered saline (PBS), followed by addition of TrypLE Express (Gibco). After incubation for 3-5 min at 37 °C, the TrypLE was inactivated by diluting with cell culture medium. The fluid volume was centrifuged at 300 RCF for 5 min. The cells were resuspended in cell culture medium for further passaging or used for patterning experiments described below. C2C12 cells between passage numbers 5 and 8 were used for patterning experiments.

### Patterning and stretching C2C12 cells in fibrin with the static stretch devices

Fibrinogen stock solutions were prepared at 50 mg mL^-1^ by dissolving 250 mg of powdered fibrinogen from bovine plasma (Sigma-Aldrick) in 5 mL of warmed DMEM. The fibrinogen solution was then filter sterilized with a 0.22 mm syringe filter. Thrombin stock solutions were prepared at 100 U mL^-1^ by dissolving 100 units of powdered thrombin from human plasma (T4393-100UN, Sigma-Aldrich) in 1 mL of warmed deionized water. Fibrinogen and thrombin stock solutions were alliquoted and stored at-20 °C until use.

After dissociating the C2C12 cells, they were pelleted and resuspended at a final concentration of 5×10^6^ cells mL^-1^ in culture medium with 5 mg mL^-1^ bovine fibrinogen and 3 U mL^-1^ thrombin. Static stretch patterning devices were assembled in a 6-well plate and 200 µL of cell-laden ECM-precursor was pipetted in the inlet and allowed to gel for 45 min at 37 °C. After gelling, the top patterning rail is removed and the transfer device is pushed into the tissue hook devices via the press-fit feature. The transfer device is then used to move the tissue hooks in tandem to the incremental stretching device in another 6-well plate. After transfer, 6 mL of growth media supplemented with 3.5 mg mL^-1^ 6-aminocaproic acid (Sigma-Aldrich) was added to each well. After 24 h in culture, tissues were randomly selected for controls (0% strain), 25% strain, and 50% strain. The tissues selected for increased strain had the tissue hooks moved from the control configuration to the appropriate strain configuration with tweezers and all tissues were changed to differentiation culture media. Differentiation culture media consisted of DMEM supplemented with 2% horse serum, 1% penicillin-streptomycin, 1X Insulin-Transferrin-Selenium (ITS-G) solution (Fisher), and 3.5 mg mL^-1^ 6-aminocaproic acid (Sigma-Aldrich). Tissues were cultured for an additional 5 days with differentiation culture media refreshed every 48 h.

### Engineered muscle tissue static stretch patch immunofluorescence staining and imaging

Engineered muscle tissue static stretch patches were rinsed twice with 1X PBS and fixed in 4% paraformaldehyde (PFA) at 4 °C for 1 h. Tissues were rinsed three times with 1X PBS before storage at 4 °C until ready for the immunofluorescence staining process. In short, the tissues were permeabilized with 0.5% Triton-X (v/v) for 1 h followed by blocking with 10% FBS for 3 h at room temperature while gently shaking on a plate shaker set at 200 RPM. Tissues were then incubated with mouse monoclonal myosin heavy chain (R&D Systems) (1:400) primary antibodies overnight at 4 °C. Tissues were then washed with a 0.2% Triton-X (v/v) solution and incubated with Alexa Fluor 488 goat anti-mouse (Jackson ImmunoResearch) (1:500) secondary antibody and Alexa Fluor 647 Phalloidin (Molecular Probe) (1:200) overnight at 4 °C. Tissues were then washed with 1X PBS and stained with 4′,6-diamidino-2-phenylindole (DAPI) (Molecular Probe) (1:1000) for 20 min at room temperature prior to imaging. Images were acquired using a 20X magnification on a Zeiss LSM 900 equipped with ZEN software. The confocal images obtained were split into individual channels per fluorophore used (MHC with Alexa Fluor 488, Phalloidin with Alexa Fluor 647, and DAPI with Alexa Fluor 405) for image analysis with fiji (imageJ) 2.16.0 or later and java 1.8.0.

### Engineered muscle tissue static stretch patch myotube alignment quantification image analysis

NIH imageJ software was used to analyze myotube alignment within the suspended tissue models to determine changes in alignment under increasing strain. Myotube alignment was quantified using two-dimensional fast Fourier transforms (2D FFT) of the confocal fluorescent images on the MYHC channel described above.^65^ 2D FFT converts an image into its frequency domain representation, revealing how pixel intensities change across the image through a magnitude plot in the frequency domain. Taking the 2D FFT of confocal images creates an image with the magnitudes of low frequencies representing background information at the center and the magnitudes of high frequencies representing edges at the border. A radius from the center is set, and pixel intensities along this radius are summed along the circular projection angles, a radial sum analysis was done by using an oval profile plugin from ImageJ.^76^ These intensities are then plotted against the corresponding angle of acquisition to obtain a plot with the representative principal myotube angle in a 360° space with respect to the direction of stretch. The peaks produced in the alignment plot represent the distribution of myotube angles within the image, more detail given in Figure S5. Aligned myotubes would display a distinct narrow peak, or a high intensity region, with the orientation of the peak indicating the dominant alignment direction of myotubes whereas not aligned myotubes would display multiple peaks or a broader peak with diffuse intensity, indicating the presence of more than one axis of alignment of myotubes.

### Engineered muscle tissue static stretch patch nuclear alignment and myotube diameter quantification image analysis

Nuclear alignment angles were measured with respect to the direction of stretch through a custom ImageJ macro on the DAPI channel of the confocal images post processing (described above). Images are split into a 10 x 10 grid overlay before the nuclei are isolated through filtering and thresholding processes. The nuclei not touching the borders of the images are then measured through ImageJ particle analysis, and the angle of their long axes with respect to the direction of stretch are recorded (Figure S3)^67,68^. Myotube diameter width was measured in ImageJ software on the MYHC channel of the confocal images post processing (described above). Three evenly spaced guidelines orthogonal to the direction of stretch were used to determine a random subset of myotubes for measurement (Figure S5)^25^. For each condition, myotubes intersecting with these guidelines were measured by a blinded analyst using standardized criteria for inclusion and exclusion, described in detail in the Supporting Information Text. Measurements were taken perpendicular to the long axis of the myotube and at three points: two points of intersection between the guideline and the edges of the myotube and a third from the center of the myotube’s intersection with the line. These measurements were then averaged to obtain the overall diameter of the myotube.

### Cell culture maintenance - 3T3 mouse fibroblasts

Initial experiments were conducted using NIH/3T3 mouse fibroblast cells obtained from ATCC. The cells were maintained in tissue culture flasks containing DMEM supplemented with 10% FBS (Gibco) and 1% penicillin-streptomycin at 37 °C, 5% CO_2_. Culture medium was changed every 48 h until cells reached 90-95% confluency, whereupon the culture was rinsed with 1X PBS, followed by addition of TrypLE Express (Gibco). After incubation for 5 min at 37 °C, the TrypLE was inactivated by diluting with cell culture medium. The fluid volume was centrifuged at 300 RCF for 5 min. The cells were resuspended in cell culture medium for further passaging or used for patterning experiments described below. 3T3 cells between passage numbers 8 and 15 were used for patterning experiments.

### Patterning 3T3 cells in collagen with two region STEAM devices

After the 3T3 cells were dissociated, two aliquots of the cells were dyed by either CellTracker Green CMFDA or CellTracker Red CMTPX at a final concentration of 25 µM in DMEM for 30 min at 37 °C. After dying, the cells were pelleted and resuspended at a final concentration of 4×10^6^ cells mL^-1^ in a liquid collagen solution. The collagen solution was prepared by mixing a stock of rat tail collagen type I (Corning) in 0.02 N acetic acid with sterile deionized water, 10X PBS, 1M HEPES buffer solution, 7.5% sodium bicarbonate solution, and 1 N NaOH to achieve a final collagen density of 5 mg mL^-1^ and kept on ice to prevent gelation. Two region STEAM devices were assembled in a 6-well plate and cell-laden ECM-precursor was pipetted in the inlet (90 µL for the inner region and 170 µL for the outer region of the 6 mm height cone-shaped device shown in Figure 5e) and allowed to gel for 45 min at 37 °C. After gelling, the top patterning rail was removed and the tissue hook device was transferred to a separate tissue holder in a separate 6- or 12-well plate. After transfer, approximately 3-4 mL of cell culture media was added to each 12-well. After 24 h in culture the tissues were fixed with 4% PFA (described above) and stored at 4 °C until imaging. Tissues were then directly transferred to a single well glass bottom plate (Cellvis) and imaged with 5X magnification on a Zeiss LSM 900 equipped with ZEN software.

## Statistical analysis

Data from the C2C12 work represents three tissues per condition. Nuclear angle alignment and 2D FFT alignment were measured from confocal images of three representative regions of interest per single tissue. All values are reported as mean ± standard deviation. One-way ANOVA with a Tukey’s post-hoc test was performed on the mean nuclear alignment angle and mean myotube diameter data using R version 4.4.2 with the dplyr and ggplot2 packages). Data visualization for Figure 4 was completed with R while Figures S3 and S7 were completed in GraphPad Prism. Comparisons with *p*-values ≤ 0.05 were considered statistically significant and denoted with an asterisk.

## Acknowledgements

This publication was supported by the National Institutes of Health (NIH) through the National Institute of General Medical Sciences award number R35GM128648 (ABT), F30HL158030 (AJH), R35GM138036 (CAD), and the National Center for Advancing Translational Sciences award number 5TL1TR002318-08 (LGB). Laura A. Milton received funding from the Fulbright Program. The content is solely the responsibility of the authors and does not necessarily represent the official views of the National Institutes of Health or other funding bodies.

## Author Contributions

JMW and AJH contributed equally to this work and they are co-first authors. JMW, AJH, ABT, and EB conceived the project with input from YCT, CAD, and NJS. ABT and EB supervised the project. JMW and AJH designed and fabricated the device with process inputs from LAK, EEB, ARV, LGB, AG, EAS, ABT, and EB. JB provided theoretical input on the flow in the two region system. JMW designed and conducted the dextran diffusion experiments with inputs from AJH, LAK, LGB, AL, and ABT. JMW designed and conducted the engineered muscle tissue experiments with inputs from AJH, LGB, LAM, ABT, and EB. JMW performed fluorescent staining and JMW, AL, MYA performed confocal imaging and processing of tissues. LAK and DAK performed image analysis with direction from JMW, AJH, ABT, and EB. JMW, AJH, EEB, ARV, LGB, AL, and EAS assisted with cell culture. EEB and EAS contributed device and hydrogel patterning photos/videos with input from JMW and AJH. JMW and AJH wrote the manuscript with significant inputs from LAK, EEB, CAD, NJS, ABT, and EB. JMW, AJH, LAK, EEB, ARV, DAK, and MYA drafted and/or generated figures. All authors have reviewed, edited, and approved of the manuscript.

