## Supplementary Information for "Suspended Tissue Engineering with Assemblable Microfluidics (STEAM)"

Affiliations
1. Department of Chemistry, University of Washington, Seattle, WA, 98195 USA
2. Medical Scientist Training Program, University of Washington School of Medicine, Seattle, WA, 98195 USA
3. School of Mechanical, Medical and Process Engineering, Queensland University of Technology, Brisbane, Australia
4. Centre for Biomedical Technologies, Queensland University of Technology, Brisbane, Australia
5. Department of Bioengineering, University of Washington, Seattle, WA, 98195 USA
6. Department of Mechanical Engineering, University of Washington, Seattle, WA, 98195 USA
7. Molecular Engineering & Sciences Institute, University of Washington, Seattle, WA, 98109 USA
8. Institute for Stem Cell & Regenerative Medicine, University of Washington, Seattle, WA, 98109 USA
9. Department of Chemical Engineering, University of Washington, Seattle, WA, 98195 USA
10. Institute for Protein Design, University of Washington, Seattle, WA, 98195 USA
11. Department of Laboratory Medicine & Pathology, University of Washington, Seattle, WA, 98195 USA
12. Department of Urology, University of Washington School of Medicine, Seattle, WA, 98195 USA

**This PDF file includes:**

Supplementary Text
Supplementary Figures
Supplementary Table
Legends for Movies

**Other Supplementary Materials for this manuscript include the following:**

Supplementary Movies

**Supplementary Text**

Inclusion and exclusion criteria for myotube diameter data analysis

In order for the diameter of a myotube to be recorded confidently by the analyst, who was blinded to the experimental conditions: the myotube must be bright enough with minimal overlap, the myotube must contain at least 2 nuclei, the myotube must be relatively cylindrical, and the myotube must intersect with the orthogonal guide lines from the ImageJ macro. For the myotube to be excluded from analysis: the myotube has a low aspect ratio (circular myosin likely indicates early stages of myogenesis), the borders of the myotube are too dark to accurately determine the diameter, the myotube edges are indistinguishable from other myotubes due to overlap, the myotube edges are disrupted by shifts in confocal tiles, or the myotubes intersecting the orthogonal guide lines are partially cut off by the borders of the confocal image.

RNA extraction, cDNA synthesis, and quantitative polymerase chain reaction

Tissue constructs were rinsed twice with 1X PBS and removed from the static stretch tissue hooks with sterile tweezers for RNA extraction. According to the manufacturer's protocol, mRNA was extracted using the Macherey-NAgel NucleoSpin RNA (Fisher Scientific). Briefly, cells were lysed in Buffer RA1 supplemented with β-mercaptoethanol, and to reduce viscosity the lysates were passed through a filter column provided with the kit. Ethanol was added to the homogenized lysate to adjust the RNA binding conditions before transfer to another filter column. A membrane desalting buffer was added to allow the rDNase digest to occur more effectively in the next step of the protocol. The DNase reaction mixture was prepared and added to the filter column, where it incubated at room temperature for 15 min. Columns were then washed with buffers to remove contaminants, followed by RNA elution in RNase-free water. RNA concentration was assessed using a BioTek Cytation5 (Agilent).

According to the manufacturer's protocol, cDNA was synthesized using the Invitrogen™ SuperScript™ IV First-Strand Synthesis System (Fisher Scientific). Briefly, 11 µL of mRNA template was mixed with 2 µL of oligioDT master mix (1 µL of oligoDT and 1 µL of dNTPs per reaction). These reactions were then incubated at 65 °C for 5 min in a thermal cycler and allowed to cool to 4 °C for at least 1 min. After incubation, 7 µL of SSIV master mix (1 µL of DTT, 1 µL of RNase inhibitor, 1 µL of SuperScript IV, and 4 µL of SuperScript IV Buffer per reaction) was added to each reaction. This reaction was then performed in a thermal cycler with the following conditions: heat to 50 °C for 10 min, then 85 °C for 10 min, then cool to 4 °C and remove to the benchtop. The resulting cDNA was diluted 1:5 in nuclease-free water for subsequent qPCR.

Quantitative polymerase chain reaction (qPCR) was done using the iTaq Universal SYBR Green Supermix (Bio-Rad) on a QuantStudio 6 Pro Real-Time PCR System (Fisher Scientific). Reactions were conducted in 10 µL volumes, consisting of 8 µL of a primer master mix and 2 µL of cDNA template. The primer master mix consisted of 2.4 µL RNase-free water, 5 µL of the SYBR Green Supermix, 0.3 uL of each primer (10 mM stock). The qPCR cycling conditions were as follows: heat to 50 °C at 1.6 °C/s and hold for 2 min, heat to 95 °C at 1.6 °C/s and hold for 10 min, followed by 40 cycles of a hold at 95 °C for an additional 15 s and decreasing the temperature to 60 °C at 1.6 °C/s and hold for 1 min. Relative gene expression levels were determined using the $2^{-\Delta\Delta Ct}$ method, with normalization to housekeeping genes. See supplementary table S1 for primer sequences.

### Supplementary Figures

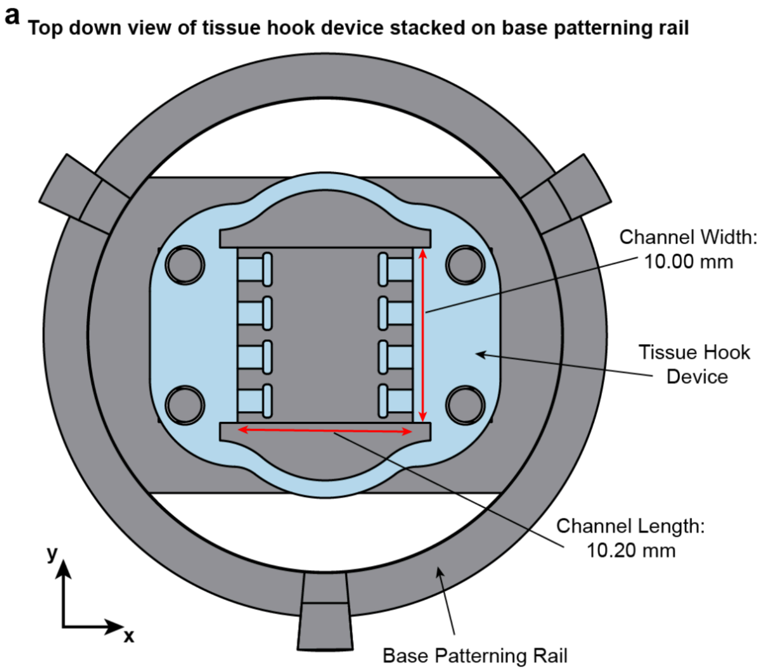


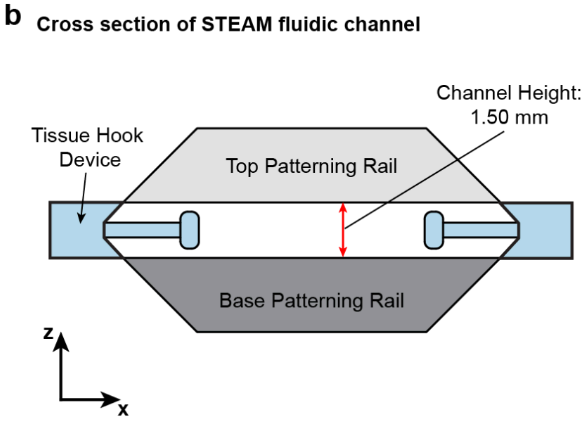


**Figure S1: Dimensions of STEAM fluidic channel used in this work**. **a** Schematic of a tissue hook device stacked on a base patterning rail (top down view). The STEAM channel length is 10.20 mm and the width is 10.00 mm. **b** Schematic of the cross section of a STEAM fluidic channel where the channel height is 1.50 mm.

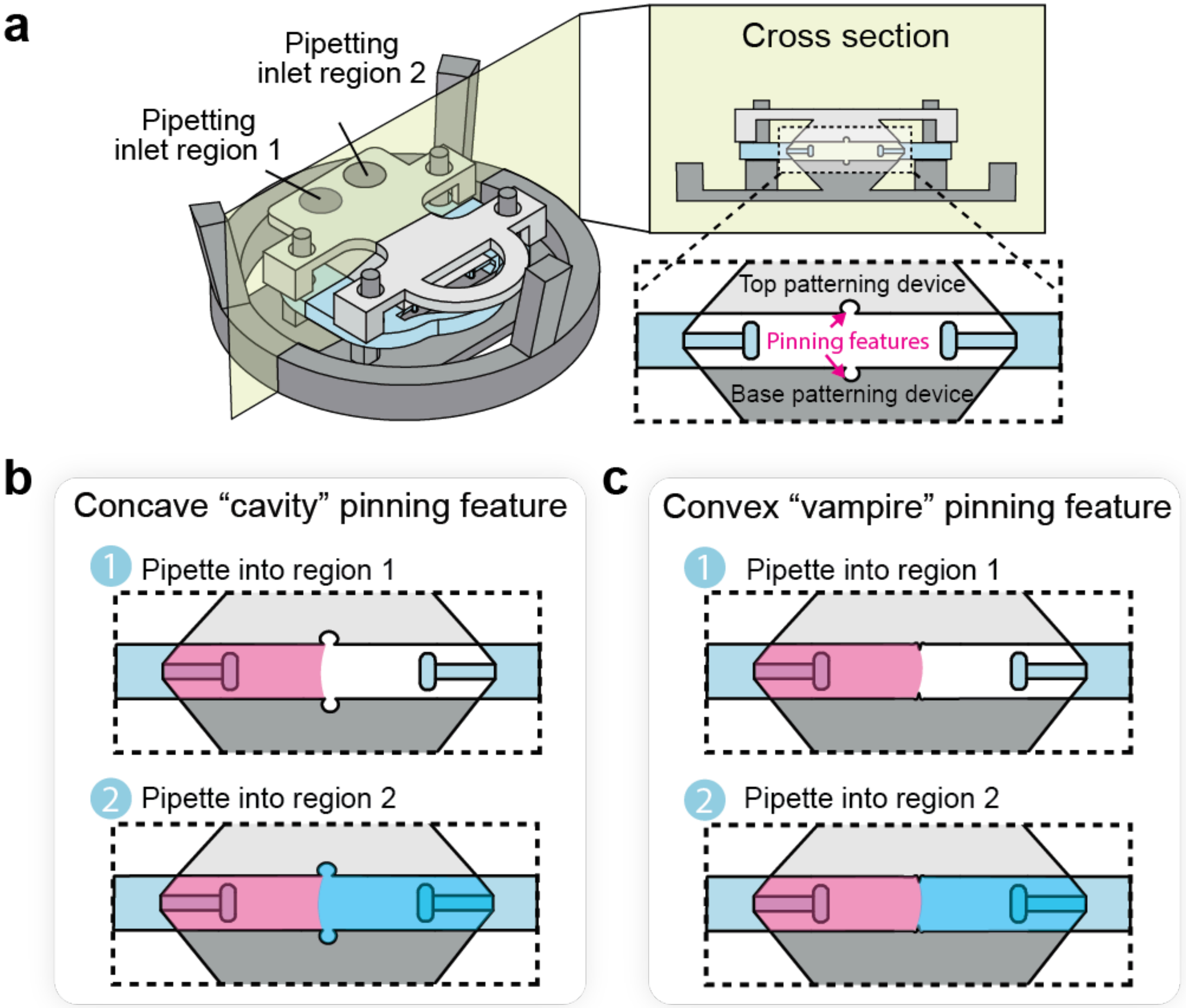


**Figure S2: Pinning features used in this work. a** Full STEAM patterning assembly with an isometric cross-sectional view of the fluidic patterning channel with pinning features integrated to the top and bottom patterning devices. **b** Concave "cavity" pinning features consist of two facing cuts into the top and base patterning devices resembling circular cavities. **c** Convex "vampire" pinning features comprise opposing reliefs from the top and base patterning devices resembling "teeth". For both pinning features, ECM-precursor is pipetted into the first region and effectively pinned to one side of the channel where a second ECM-recursor can then be pipetted into the second region to form a contiguous tissue.

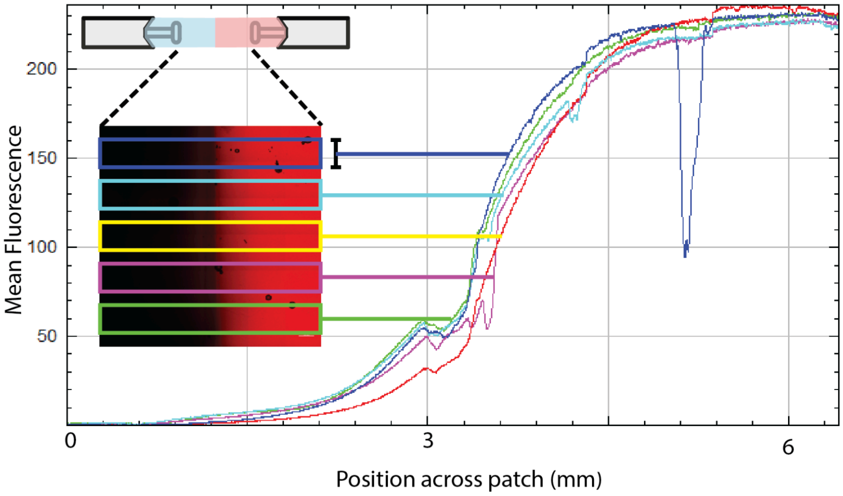


**Figure S3: Workflow for dextran diffusion fluorescent quantification from confocal images.** A maximum projection confocal image is analyzed by overlaying five evenly spaced rectangular regions perpendicular to the direction of patterning with an example shown above. The fluorescent intensity at each point was then averaged across the width of each rectangular region producing a line. These lines are averaged and plotted to determine the overall average fluorescence profiles of dextran per patch as seen in Figure 3 and Figure S4.

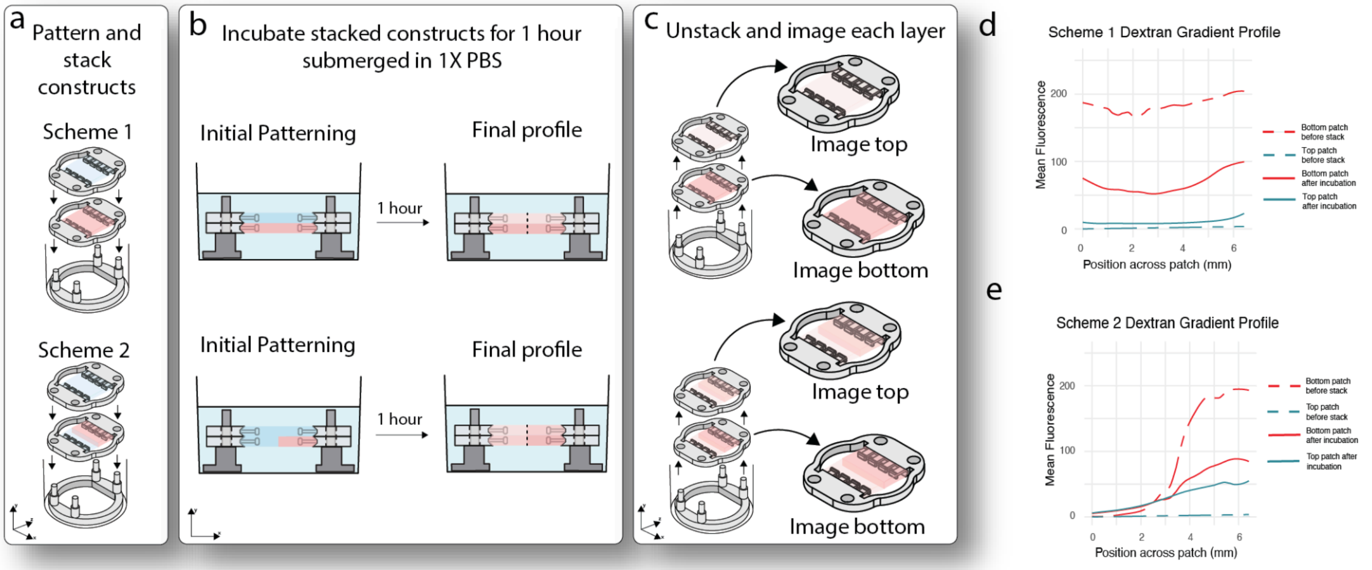


**Figure S4: Additional replicate data of dextran diffusion model.** **a** STEAM planar patch constructs were patterned and stacked on top of each other in a 12 well plate. Experimental scheme 1 was a dextran-infused single region collagen patch with a blank collagen patch on top while experimental scheme 2 was a two region patch with dextran-infused collagen on one side and blank collagen on the other with a blank collagen patch on top. **b** Stacked constructs are then submerged in 1X PBS for 1 h to allow for diffusion of dextran. **c** The constructs were then unstacked and confocal imaged separately. **d** Fluorescent profiles were generated from intensity of dextran of the collagen patches before incubation (dashed line) and after incubation (solid line), experimental scheme 1 revealed diffusion with an increase of dextran in the top collagen patch (blue) and a decrease in the bottom collagen patch (red). **e** For experimental scheme 2, the same trend as scheme 1 was revealed with an increase of dextran in the top collagen patch (red), however, a spatial gradient was detected due to the initial two region patterning of the bottom collagen patch (blue) and initial placement of dextran.

a

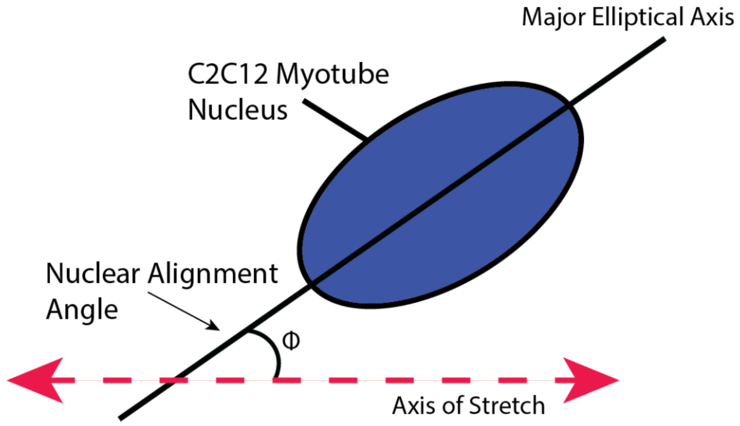


b

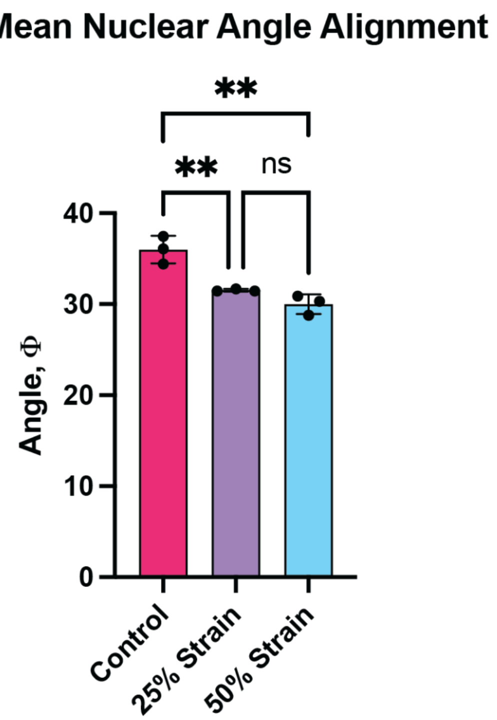


**Figure S5: Nuclear alignment is increased with strain. a.** Nuclear alignment was analyzed where confocal images of the DAPI channel were split into a 10x10 grid. Image processing was done to each sub-image to remove internal and external noise within the image. A threshold and watershed function from ImageJ was then applied to isolate individual nuclei. An analyze particles function (ImageJ) was done to superimpose ellipses on top of the nuclei to acquire the major elliptical axis. The angle, Φ, was then measured with respect to the axis of stretch as shown with this diagram. **b.** Nuclear alignment analysis was completed and the mean nuclear angle was compared. The 25% strain and 50% strain conditions were significantly different when compared to the control condition, however the 25% and 50% conditions were not significant when compared to each other ($p \geq 0.05$). Each data point is the mean nuclear alignment angle from a separate tissue, three tissues per condition for a single experiment. Error bars represent the mean ± standard deviation. Statistical analysis was performed with One-way ANOVA with a Tukey's post-hoc test where **$p \leq 0.01$ and *ns* is not significant ($p > 0.05$).

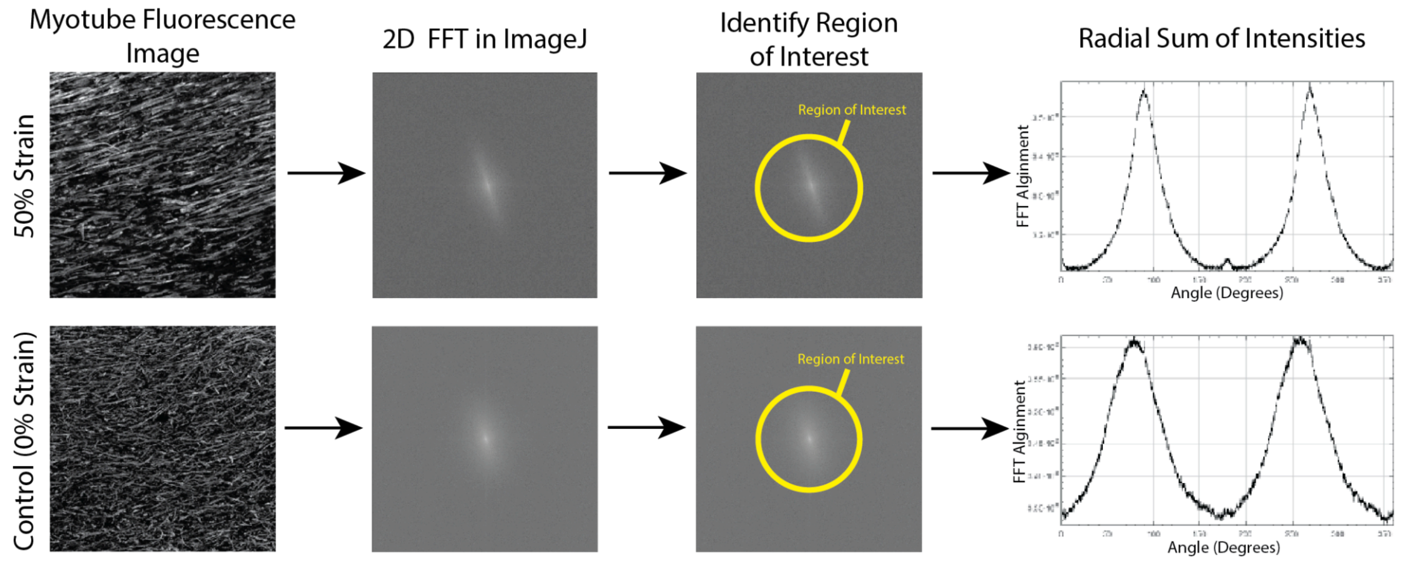


**Figure S6: Workflow from a fluorescence image to obtain a 2D FFT alignment plot.** First, tissues were immunostained with myosin heavy chain antibody to visualize myotubes and imaged with confocal microscopy. 2D FFT images were then obtained using an NIH ImageJ macro. 2D FFT images represent information pertaining to a defined frequency space domain that is converted from the corresponding real space domain of the confocal image. This frequency domain can subsequently be used to observe the rate of change of pixel intensity distribution throughout an image. This rate of change of pixel intensities can be plotted around the origin in a manner to represent the degree of alignment within the image. The 2D FFT image is thus represented where low-frequency signals and the background are placed at the center, and the high-frequency signals, which represent edges and noise, are present in the peripheral regions with reference to the origin. Therefore, a defined region of interest at a set pixel radius from the center which is kept consistent across each image. The region of interest pixel intensities are summed up radially along the circular projection angles by using an oval profile plugin from ImageJ. These intensities are then plotted against the corresponding angle of acquisition to obtain the final 2D FFT alignment plot for a fluorescence image to show a representative principal angle of myotube alignment in a 360° space. Height and overall shape of the peak represent the degree of alignment of myotubes within the image. A high and narrow peak in the plot represents a uniform degree of alignment while a broad peak indicates the presence of more than one axis of alignment of myotubes. For a full random orientation, no recognizable peak would be observed. Note: The angle of the FFT peak corresponds to the direction perpendicular to the fiber orientation. Figure recreated from Vajanthri et al. using their workflow with images and data from our own experiments.

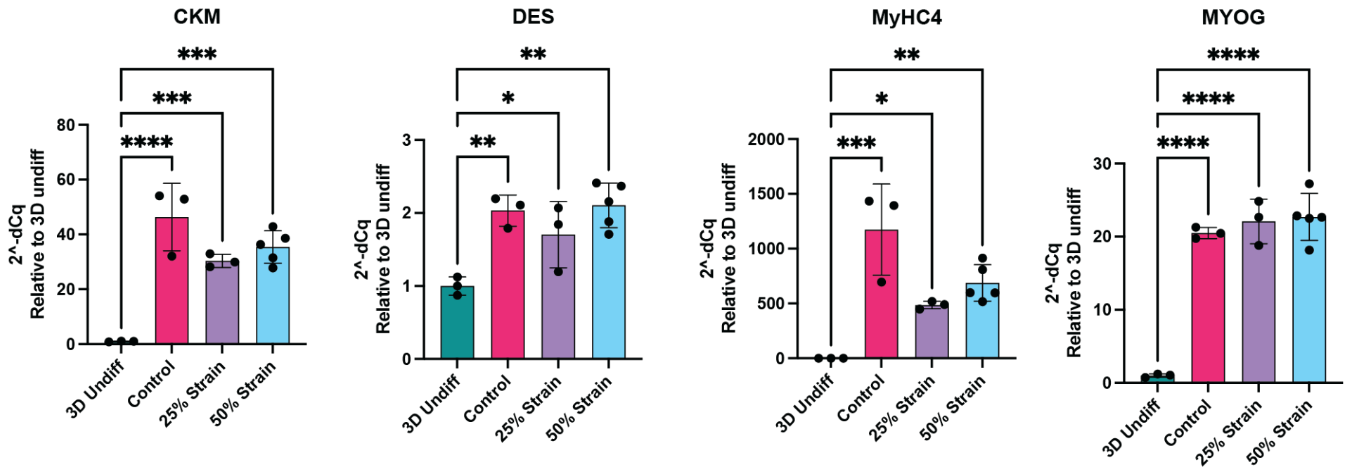


**Figure S7: mRNA expression of muscle differentiation markers compared to undifferentiated 3D constructs.** Muscle differentiation markers, such as *MyHC4*, *MYOG*, *DES*, and *CKM*, were chosen to confirm successful differentiation from muscle myoblasts to myotubes when compared to 3D undifferentiated muscle tissues using STEAM devices. Relative gene expression levels were determined using the $2^{-\Delta\Delta Cq}$ method, with normalization to housekeeping genes (*GAPDH* and *PGK*). Error bars represent the mean ± standard deviation. Statistical analysis was performed with One-way ANOVA with a Brown-Forsythe post-hoc test. Significant upregulation was determined, *$p \leq 0.05$, **$p \leq 0.01$, ***$p \leq 0.001$, ****$p \leq 0.0001$).

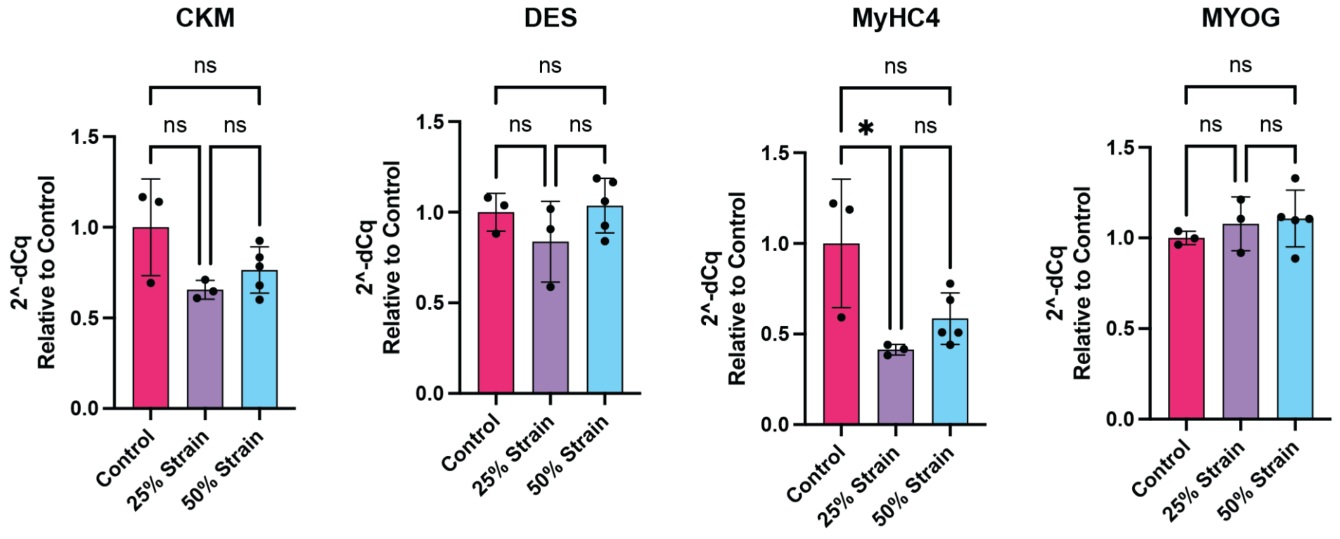


**Figure S8: mRNA expression of muscle differentiation markers compared to static stretch control.** Muscle differentiation markers, such as *MyHC4*, *MYOG*, *DES*, and *CKM*, were chosen to determine mRNA expression levels when compared within the 3D differentiated control (no strain) when STEAM devices are used. Notably, there is no marked difference between conditions. Relative gene expression levels were determined using the $2^{-\Delta\Delta Cq}$ method, with normalization to housekeeping genes (*GAPDH* and *PGK*). Error bars represent the mean ± standard deviation. Statistical analysis was performed with One-way ANOVA with a Brown-Forsythe post-hoc test where *$p \leq 0.05$ and *ns* is not significant ($p > 0.05$).

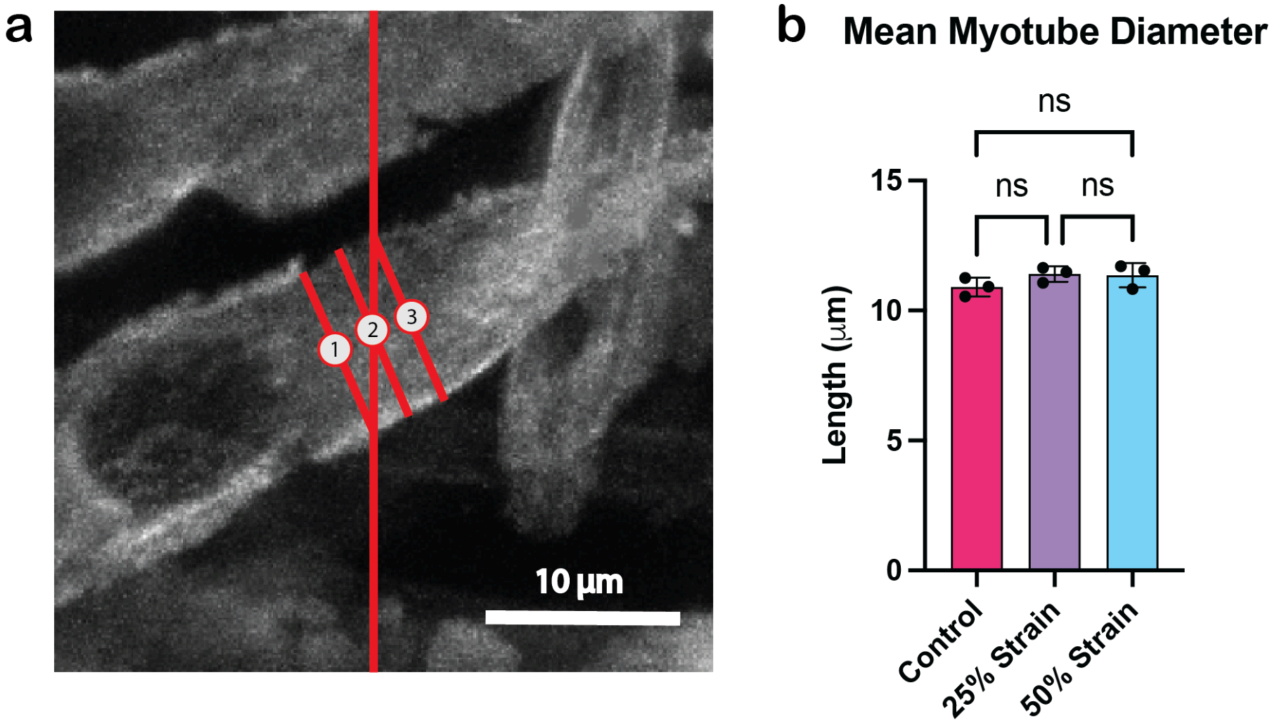


**Figure S9: Myotube diameter is consistent across all levels of strain. a.** Myotube diameter was measured with a NIH “3 LINES.ijm” ImageJ macro to create three evenly spaced reference lines orthogonal to the direction of stretch for each image. Myotubes overlapping the red reference lines were sampled with inclusion and exclusion criteria described above. An average of three diameter measurements, labeled “1”, “2”, and “3” in the example image, were taken for each myotube recorded. **b.** The mean myotube diameter for each condition was then compared. Each data point is the mean myotube diameter from a separate tissue, three tissues per condition for a single experiment. Error bars represent the mean ± standard deviation. Statistical analysis was performed with One-way ANOVA with a Tukey's post-hoc test. No statistically significant differences were found ($p > 0.05$).

**Table S1: Primer Sequences**

| Target | Species | Forward (5’-3’) | Reverse (5’3’) |
|---|---|---|---|
| MYOG | Mouse | AGTACATTGAGCGCCTAC | CAAATGATCTCCTGGGTTG |
| MYHC4 | Mouse | TCAAATTATCAGTGCCAACC | ACTTCTCTAGCAGATAGGTTTC |
| DES | Mouse | CAGGATCAACCTTCCTATCC | CTGTCTTTTTGGTATGGACTTC |
| CKM | Mouse | ACAAAAGCTTCCTTGTGTG | AGATCTCCTCAATCTTCTGC |
| GAPDH | Mouse | ACACCCTCAAGATTGTCAGCAA | TCATAAGTCCCTCCACGATGC |
| PGK | Mouse | CAAAATGTCGTCTTCCAACAAG | AACGTTGAAGTCCACCCTCAT |

**Movie S1 (separate file).** Example of patterning a single region planar patch tissue with STEAM. Blue colored agarose is pipetted into the inlet of the top patterning rail and left to gel.
**Movie S2 (separate file).** Example of removal process of a single region planar patch tissue with STEAM. Tweezers are used to remove the top patterning rail first, then the overall tissue hook device with a suspended tissue in blue colored agarose is lifted from the base patterning rail.
**Movie S3 (separate file).** Example of patterning a two region planar patch tissue with STEAM. Blue colored agarose is pipetted into one inlet of the top patterning rail followed by pink colored agarose in the other inlet and left to gel.
**Movie S4 (separate file).** Example of removal process of a two region planar patch tissue with STEAM. Tweezers are used to remove the top patterning rail first, then the overall tissue hook device with a two region suspended tissue in blue and pink colored agarose is lifted from the base patterning rail.
**Movie S5 (separate file).** Demonstration of five wave-shaped agarose constructs assembled with colored food dye diffusing over 2 hours to create a gradient vertically through the stack and laterally along the wave contours. The top layer was colored green and the bottom layer was colored purple, with three clear layers stacked in between. This time lapse video was created from 2 h of raw video footage and is 82.76% faster than the original real time speed. At the end of the 2 h of diffusion, a photo was taken for the image in Figure 5g. Scale bar is 1 cm.
